# *Vibrio campbellii* chitoporin: thermostability study and implications for the development of therapeutic agents against *Vibrio* infections

**DOI:** 10.1101/2020.06.24.169508

**Authors:** Anuwat Aunkham, Albert Schulte, Wei Chung Sim, Watcharin Chumjan, Wipa Suginta

## Abstract

*Vibrio campbellii* (formerly *Vibrio harveyi*) is a bacterial pathogen that causes vibriosis, which devastates fisheries and aquaculture worldwide. *V. campbellii* expresses chitinolytic enzymes and chitin binding/transport proteins, which serve as excellent targets for antimicrobial agent development. We previously characterized *Vh*ChiP, a chitooligosaccharide-specific porin from the outer membrane of *V. campbellii* BAA-1116. This study employed far-UV circular dichroism and tryptophan fluorescence spectroscopy, together with single channel electrophysiology to demonstrate that the strong binding of chitoligosaccharides enhanced thermal stability of *Vh*ChiP. The alanine substitution of Trp^136^ at the center of the affinity sites caused a marked decrease in the binding affinity and decreased the thermal stability of *Vh*ChiP. Tryptophan fluorescence titrations over a range of temperatures showed greater free-energy changes on ligand binding (Δ*G*°_binding_) with increasing chain length of the chitooligosaccharides. Our findings suggest the possibility of designing stable channel-blockers, using sugar-based analogs that serve as antimicrobial agents, active against *Vibrio* infection.

## Introduction

Chitin is a natural, insoluble polysaccharide composed of repeating units of *N*-acetylglucosamine (GlcNAc) linked by *β*-1,4-glycosidic linkages, and is primarily found in the cell walls of fungi and the exoskeletons of arthropods and crustaceans. Global demand for seafoods has promoted shellfish cultivation worldwide, with the total world crab production, for instance, being over 200 million tons in 2016 as reported by World Food Program (WFP) (https://www.wfp.org/). Since a large portion of the weight of a crustacean is inedible, the shellfish farming and processing industry produces approximately 6 – 8 million tons of waste shells worldwide annually, of which 1.5 million is contributed by Southeast Asia. These waste shells, which contain 20 – 58% chitin, are often simply discarded, either in the ocean or into landfills. Although chitin is biodegradable, its natural components are refractory to degradation by naturally occurring enzymes, which has led to its accumulation over time and poses a serious threat to coastal ecosystems [1,2,3,4]. Chitin has received much attention in recent years from entrepreneurs and researchers who are working on eco-friendly technology for the development of renewable bioenergy, biomedicine and biotechnology. Chitin serves as a suitable biomaterial because of its low cost but high availability as waste biomass and its excellent biocompatibility. Chitin is therefore regarded as a sustainable future energy source and an alternative to depleting non-renewable fossil fuel resources. Previous reports showed that the chitinolytic bacteria *Aeromonas hydrophila* and *Bacillus circulans* can be used in microbial fuel cells for electricity generation. [5,6]. In other applications, accelerating the biodegradation of chitin will not only reduce pollution but also produce high-value products [7], as chitin wastes can be conveniently degraded by endochitinases and exo-*N*-acetylglucosaminidases, generating bioactive chitooligosaccharides of high pharmacological/biotechnological value [8,9].

*Vibrio* species are the most abundant bacteria in aquatic habitats, and of these *Vibrio campbellii* (formerly *V. harveyi*) is a major member. The bacteria are known to cause luminous vibriosis, which destroys commercially farmed shrimps and other aquaculture and is a cause of global economic loss [10, 11, 12, 13]. *V. campbellii* use chitin for energy production through the chitin catabolic pathway, in which chitin is degraded first to chitooligosaccharides of various lengths, but mainly to chitobiose, by chitinases such as ChiA [14,15]. Small sugar molecules, such as *N*-acetylglucosamine and (GlcNAc)_2_, may enter bacterial cells through general diffusion porins, while longer chitooligosaccharides are transported through a chitooligosaccharide-specific channel, known as chitoporin or ChiP [16,17]. Once within the periplasmic compartment, these oligosaccharides are degraded by exo-*N*-acetylglucosaminidase (GlcNAcase) [18], yielding GlcNAc and (GlcNAc)_2_, which are transferred by chitin-binding protein (CBP) to a (GlcNAc)_2_-specific ABC transporter or to GlcNAc-specific PTS transporters and transported through the inner membrane into the cytoplasm, where they are catabolized and used as primary carbon and nitrogen sources [19, 20, 21]. Although the operation of the chitin catabolic cascade of marine *Vibrio sp*. is partially understood, it is known to be associated with several chitin-inducible genes in the *chib* operon that are controlled by a two-component membrane-bound histidine kinase, identified as chitin sensor (ChiS) [22]. Chitin degradation and transport of chitin degradation products are crucial for the *Vibrio* bacteria to survive and thrive in aquatic environments that are chitin-enriched but glucose-deficient. *Vh*ChiP from *V. campbellii* strain ABB-1116 was shown to form a stable trimeric channel with an average conductance of 1.8 ± 0.3 nS in artificial phospholipid membranes [17]. Notwithstanding its ability to transport a range of chitooligosaccharides, the kinetic parameters based on time-resolved single-channel recordings suggested that *Vh*ChiP preferred long-chain substrates and the binding constant (*K*) of 0.5 × 10^6^ M^−1^ for chitohexaose was the greatest among the chitooligosaccharides tested [17, 23]. The crystal structures of *Vh*ChiP in the absence and presence of chitotetraose and chitohexaose were reported recently [24]. The overall structure is a trimeric assembly, each subunit containing 16 *β*-strands connected by 8 extracellular loops and 8 periplasmic turns. The longest loop (L3) was found to protrude into the protein pore and contains a prominent Trp residue (Trp^136^) at the center of the elongated affinity site. The *Vh*ChiP structure with bound chitooligosaccharides shows that the sugar passage is highly asymmetric, one side being more hydrophobic while the other contains more charged residues. The *N*-termini of the trimeric pores each contain a helix of 9 amino acids (the so-called *N*-plug), the role of these *N*-plugs being to regulate the opening and closing of the channel. In this present study, the effects of chitooligosaccharide binding on the structural integrity of *Vh*ChiP were investigated using circular dichroism and fluorescence spectroscopy. The findings offer a strategic concept for the development of potential anti-*Vibrio* agents based on chemoenzymatic synthesis of chitooligosaccharide analogues that can form a stable complex with the *Vh*ChiP channel.

## Results

### Structural dissection of chitooligosaccharide-*Vh*ChiP interactions

The crystal structures of *Vh*ChiP WT in complex with chitohexaose (pdb id 5MDR) and chitotetraose (pdb id 5MDS) were published recently [24]. Figure 1 shows the structural details of *Vh*ChiP, focusing on the affinity sites that are distributed along the sugar passage, where a chitooligosaccharide binds and is subsequently translocated. **Figure 1A** depicts the trimeric assembly of *Vh*ChiP in the outer membrane of the bacterium, with two monomers (**red and blue**) shown in surface representation, while the third monomer is shown as a cartoon representation (**green**). In the absence of the sugar substrate, the pore-restricting loop L3 (**orange**), containing two short helices, is found to protrude into the middle of the protein pore and interact with the *N*-helical segment of 9 amino acids (the so-called *N*-plug) from the neighboring monomer (**a red cylinder**). In the open state, a chitohexaose chain (**shown in a stick model of one representative pore**) fully occupies the channel lumen and replaces the interactions between loop L3 and the *N*-plug, causing the *N*-plug to be ejected from the pore to a position in the periplasmic space. The distance of the *N*-plug between the closed and open states of the channel is measured from the *C*_α_ of the first amino acid residue (labelled *C*_α_-Asp^1^) to be 35.7 Å (**Fig. 1B**).

**Fig. 1.**
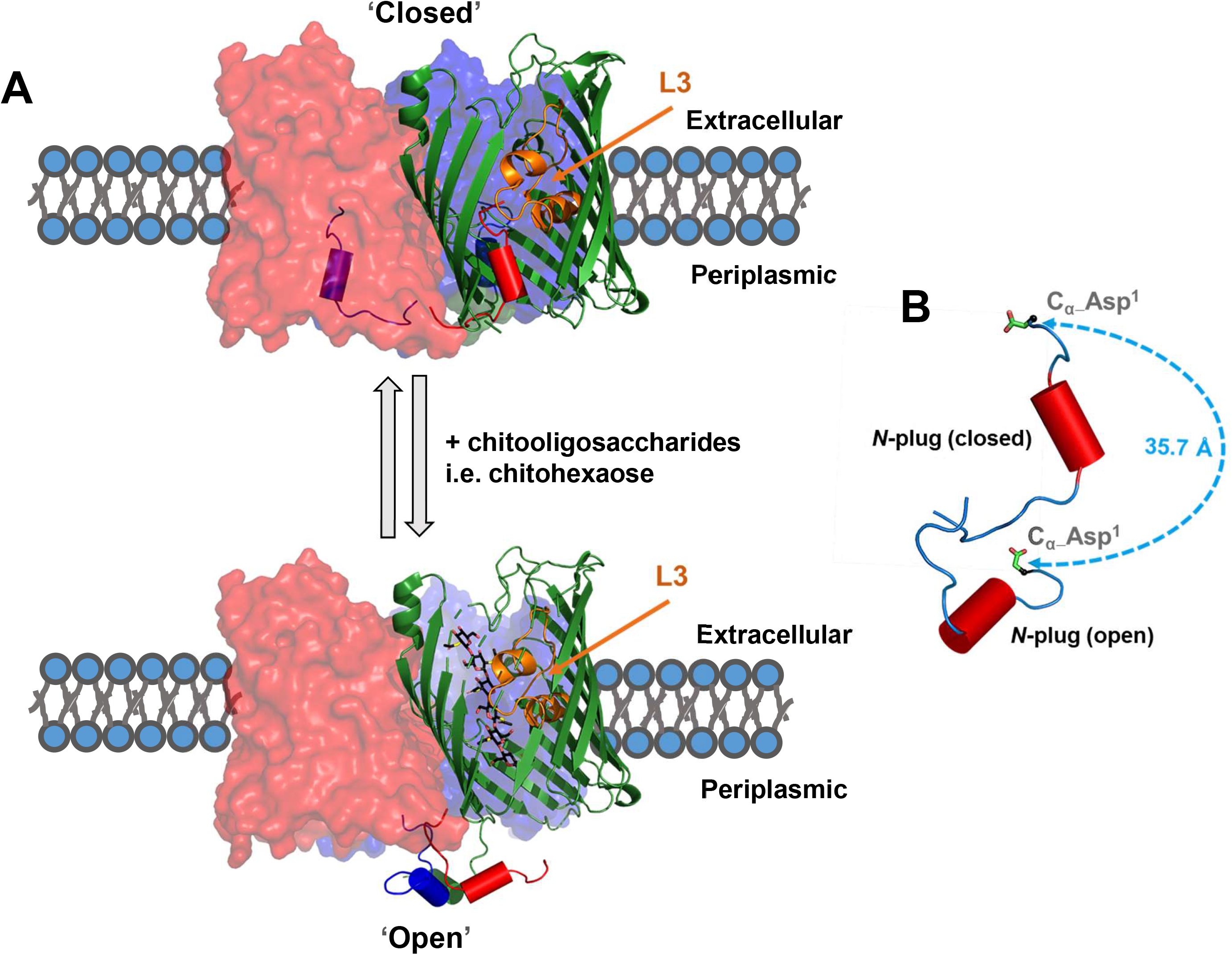
Structural dynamics of closed (PDB code **5MDQ**) and open states of the *Vh*ChiP porin (PDB code **5MDR**). **(A)** Two subunits (red and blue) of trimeric *Vh*ChiP are shown as surface representations, while the third subunit (green) is shown as a cartoon representation. The N-plug of each subunit is shown as cylinder. Loop L3 is shown in orange. Chitohexaose is shown as sticks and colored black for C, red for O, and yellow for N. Dashed blue circles are residues that interact with chitohexaose, but not chitotetraose. (**B**) A zoom in of a representative *N*-plug between the open and closed states. The distance of the plug between the two states of *Vh*ChiP is measured from the C_α_ atom of Asp^1^.

Tight substrate-binding affinity was observed in titration experiments, which showed that *Vh*ChiP has greater affinity for chitohexaose than for chitotetraose, and this can be conveniently elaborated by LIGPLOT analysis, which shows the detail of interactions between the channel and chitohexaose (**Fig. 2A**) compared with chitotetraose (**Fig. 2B)**. Both sugars are seen in extended conformations, with chitohexaose stretched fully along the channel lumen and the six pyranose rings aligned in an alternating fashion, making extensive interactions with the affinity sites +1, +2, +3, +4, +5 and +6. The two outermost sugar rings, at affinity sites +5 and +6, are surrounded by the hydrophobic sidechains of Trp^136^ and Trp^331^ on one side and of Phe^84^ on the other, while the innermost sugar ring stacks against the hydrophobic sidechain of Trp^123^ at affinity site +1. The C2-acetamido groups of each sugar ring of both chitooligosaccharides make relatively strong hydrogen bonds either with nearby charged residues or with water molecules within 3.5 Å. Hydrogen-bond contacts of the sugar moieties are formed with the surface residues Asp^53^, Arg^94^, Asp^122^, Asn^127^, Asp^135^, Arg^148^, Asn^336^ and Glu^347^. Note that the highly charged residues form H-bonds (**red dashed lines**) with the C2-acetamido and -OH groups in GlcNAc units at internal affinity sites. In contrast, the chain of chitotetraose was clearly bound at the high-affinity sites +2 to +5 (**Fig. 2B**), interacting with the same residues that bind chitohexaose at those affinity sites. The most notable difference is that chitotetraose lacks interactions with Asn^336^ and Trp^123^ at the +1 site and Trp^331^ at the +6 site (**residues with cyan dashed circles**); these residues make direct contacts with chitohexaose. With close inspection of the electron-density map of chitotetraose, we detected sporadic electron density at the affinity sites +1 and +6, which may indicate low occupancy of the chitotetraose chain in the alternative binding modes +1 to +4 or +3 to +6. However, we did not model the GlcNAc rings in these sites due to inadequate electron density.

**Fig. 2.**
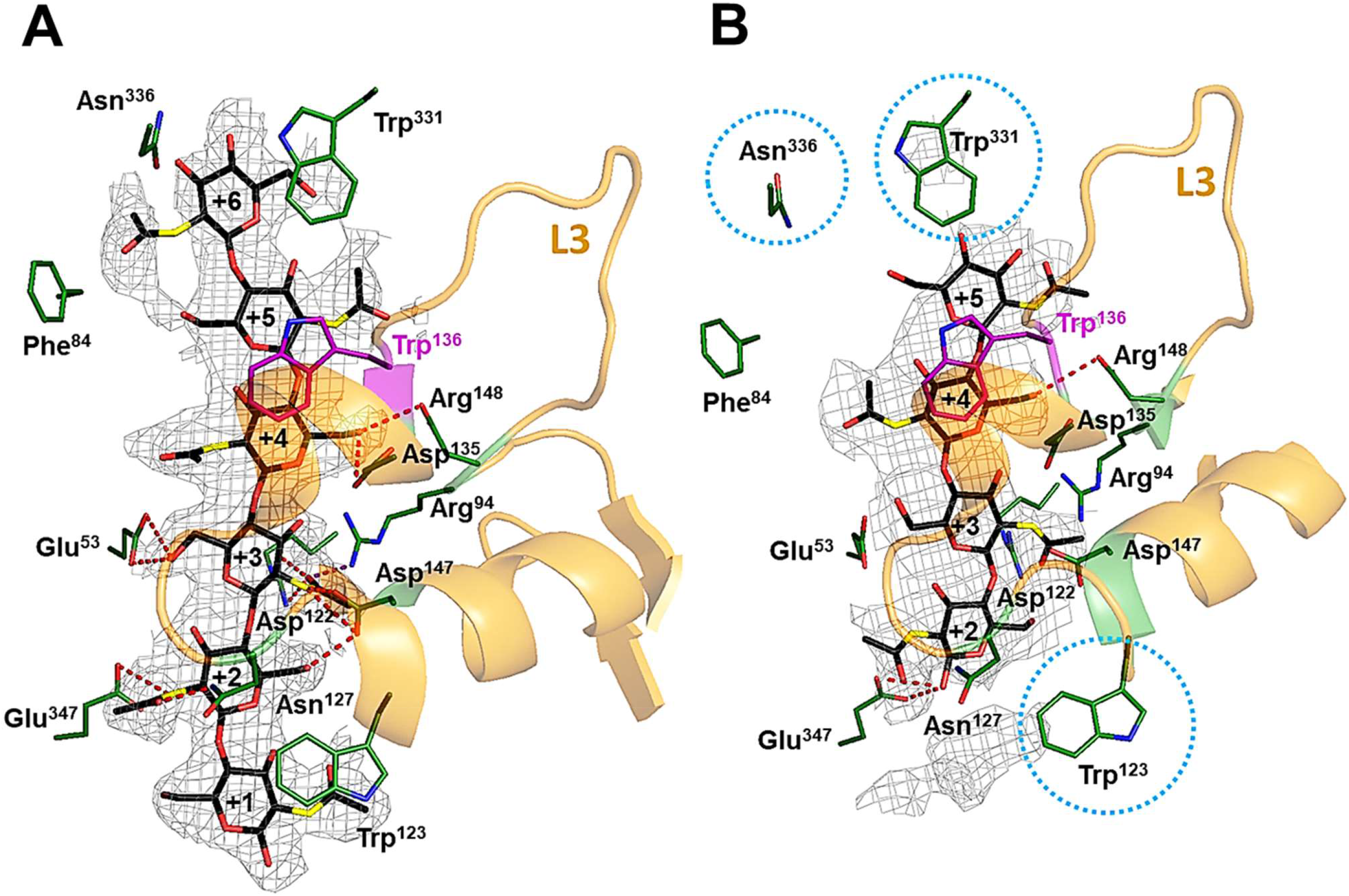
LigPlot analysis of *Vh*ChiP binding to chitooligosaccharides. (**A**) chitohexaose (PDB code **5MDR**) and (**B**) chitotetraose (PDB code **5MDS**). The affinity sites are designated +1 to +6. Sugar-binding residues are shown as sticks and colored by atom, green for carbon, blue for nitrogen and red for oxygen. Hydrogen bonds are shown as red dashed lines. Sugar backbones are shown as sticks with atoms colored black for C, red for O and yellow for N. The 2*F*_o_-*F*_c_ omit maps of both chitooligosaccharides are contoured at σ = 2.0. Cyan dashed circles are the residues that interact with chitohexaose, but not with chitotetraose. Hydrogen bonds are set at 3.0 Å.

### Temperature-induced subunit dissociation

We previously demonstrated by SDS-PAGE analysis that the trimeric *Vh*ChiP remained intact when subjected to increasing temperature, beginning to dissociate only when the temperature was raised above 60 °C [17]. In this set of experiments, we analysed temperature-induced subunit dissociation in the absence and presence of two representative chitooligosaccharides, chitohexaose and chitotetraose. The purified protein, either with or without the chitosugars, was heated for 10 min with 10 °C temperature increments (at 30, 40, 50, 60, 70, 80, 90, and 100 °C), and the protein was then examined by SDS-PAGE. **Figure 3** shows the effects of temperature treatment on the migration of the WT *Vh*ChiP (**Fig. 3A**) and the W136A mutant (**Fig. 3B**) on SDS-PAGE gels. In the absence of chitooligosaccharide, WT migrated to just below the position of the 75-kDa standard when exposed to 30-50 °C. This protein band is presumed to be the trimeric *Vh*ChiP (**Fig 3A**, **top left panel**). Note that we have shown previously that in the case of *E. coli* porin (*Ec*ChiP), the unheated porin migrated faster than the heat-denatured protein on SDS-PAGE and the faint bands of the folded protein were observed, since they were not surface accessible, and thus could not bind strongly to the Coomassie dye [25]. When *Vh*ChiP was heated at 60 °C, two protein bands appeared, the faint high-molecular weight band of the intact protein and a band representing the dissociated protein at about 40 kDa. The latter band represents the monomeric form produced by dissociation of the three subunits. Complete dissociation of the protein trimers to monomers was seen at temperatures of 70 °C and above. The subunit migration profile of *Vh*ChiP WT in the presence of the non-binding oligosaccharide maltohexaose resembled that of the *Vh*ChiP without chitooligosaccharide (**Fig 3A**, **top right panel**). In contrast, the protein bound to chitotetraose was almost completely dissociated to monomers at ≥ 70 °C (**Fig. 3A**, **bottom left panel**), while the *Vh*ChiP complexed with chitohexaose showed more resistance to heat treatment, beginning to dissociate to monomers at 70 °C and completely dissociating only when the temperature was raised to 80 °C and above (**Fig. 3A**, **bottom right panel**). Mutation of the critical residue Trp^136^ on loop L3 to Ala (mutant W136A) clearly destabilized the protein trimers. As shown in **Fig. 3B** (**top left panel)**, the mutated *Vh*ChiP already began to dissociate at 50 °C and was fully dissociated at 60 °C. Addition of maltohexaose (**Fig. 3B**, **top right panel**) or chitotetraose (**Fig. 3B**, **bottom left panel**) did not change the protein’s heat susceptibility, as the dissociation profile of the W136A mutant was identical to that without sugar added. However, chitohexaose clearly restored the protein’s stability, shifting the dissociation temperature to 70 °C (**Fig. 3B**, **bottom right panel**).

**Fig. 3.**
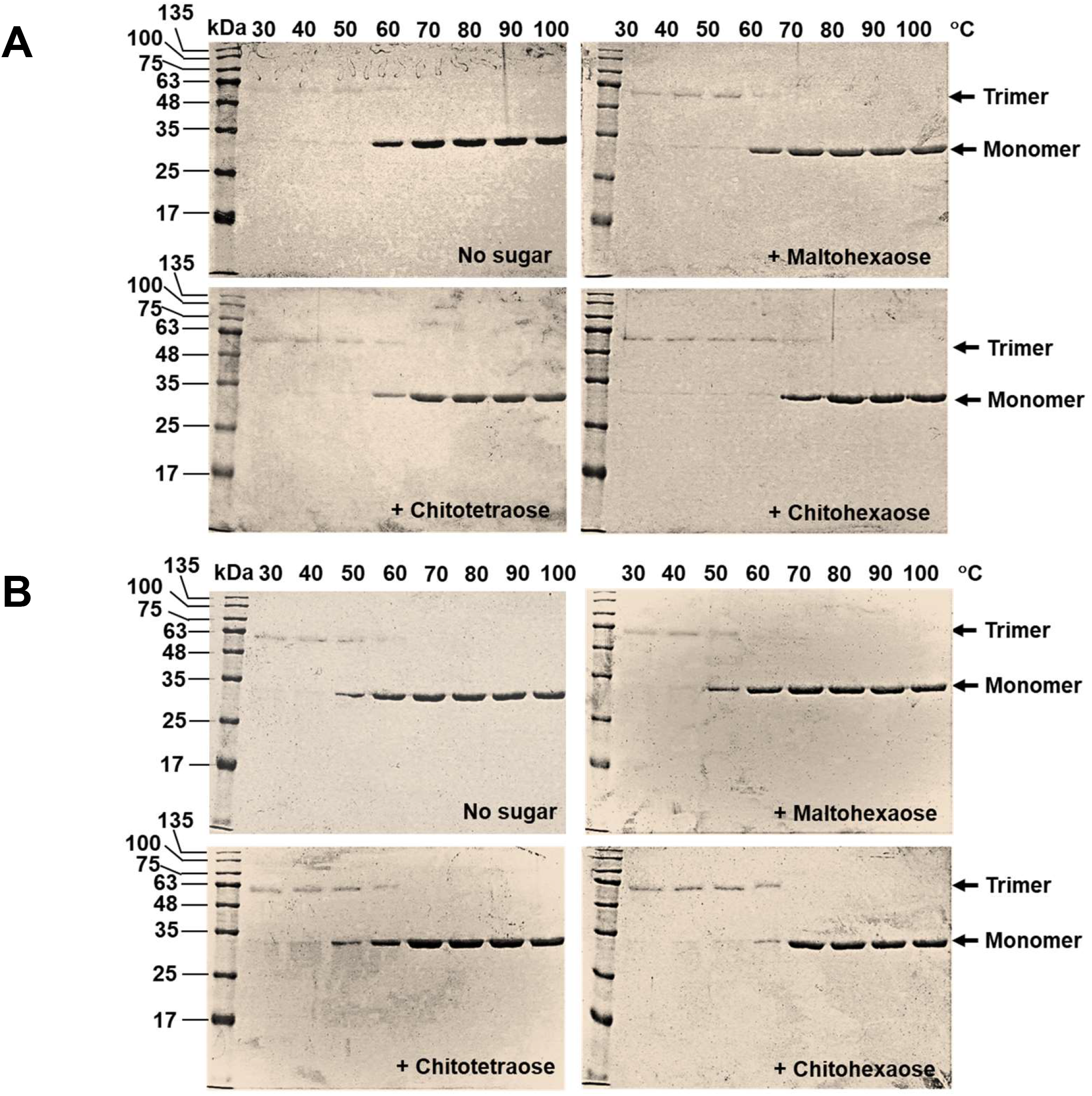
Temperature-induced subunit dissociation of *Vh*ChiP. Mixtures of purified *Vh*ChiP (10 μM) and chitooligosaccharide (100 μM) were incubated at different temperatures for 10 min and then analysed by 10% SDS-PAGE, followed by Coomassie Blue staining. (**A**) WT and (**B**) W136A. Panels: top left, *Vh*ChiP with no sugar; top right, *Vh*ChiP with maltohexaose; bottom left, *Vh*ChiP with chitotetraose; and bottom right, *Vh*ChiP with chitohexaose. Heat treatment: lane 2, 30 °C; 3, 40 °C; 4, 50 °C; 5, 60 °C; 6, 70 °C; 7, 80 °C; 8, 90 °C; 9, 100 °C. The folded protein is labelled as a trimer, while the unfolded protein is labelled as a monomer.

Gel filtration chromatography was used to confirm that the two protein bands shown on SDS-PAGE gels are the trimeric and monomeric states of *Vh*ChiP. **Figure 4A** shows a Superdex^®^ 200 chromatographic profile of four standard proteins of known molecular weight (**grey dotted lines**) and the *Vh*ChiP samples (**black lines**) treated at 30 and 100 °C. <u>*Vh*</u>ChiP WT treated at 30 °C was eluted at a position between bovine serum albumin (66 kDa) and glucose oxidase (160 kDa), the apparent molecular weight of the protein, estimated from its distribution coefficient (*K*_av_), being 125 kDa, consistent with the trimeric form of *Vh*ChiP (**Fig. 4B**). On the other hand, the protein treated at 100 °C was eluted at a position between lysozyme (14 kDa) and ovalbumin (43 kDa), its apparent molecular weight of 38 kDa being consistent with a monomeric structure.

**Fig. 4.**
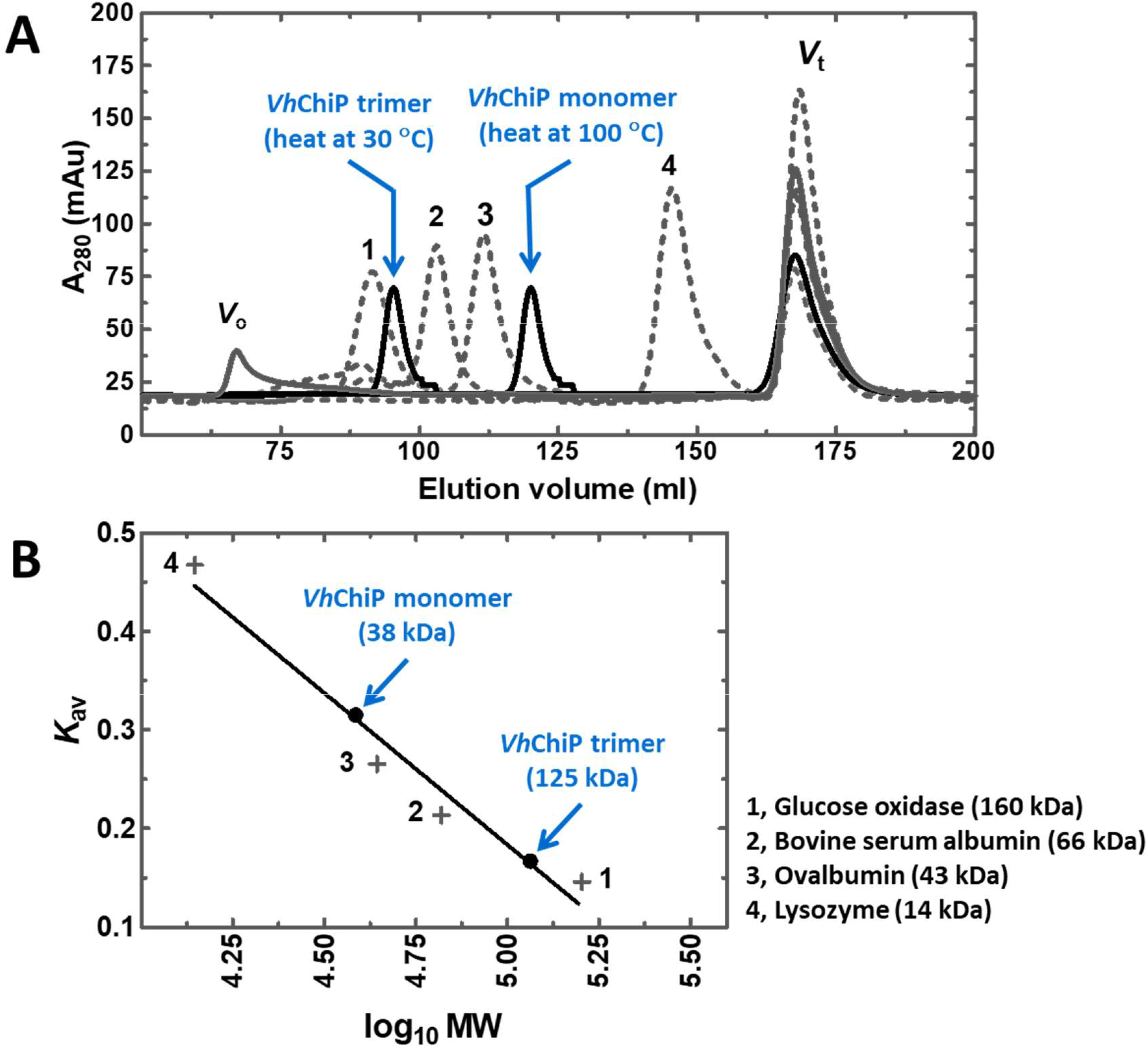
Molecular weight determination of trimeric and monomeric states of *Vh*ChiP. (**A**) *Vh*ChiP (3.0 mg) was chromatographed on a Superdex S-200 column. Standard proteins applied were: (1) glucose oxidase, (2) bovine serum albumin, (3) ovalbumin and (4) lysozyme. Blue dextran was used to obtain the void volume (V_o_) and DNP-lysine the total bed volume (V_t_) of the column. (**B**) The distribution coefficient (*K*_av_, estimated from Eq. 1) plotted against the logarithmic molecular weights of standard proteins. The standard curve was constructed from a linear curve fit available in GraphPad Prism *v*. 6.0. The molecular weights of *Vh*ChiP pre-treated at 30 or 100 °C were determined from their corresponding *K*_av_ values as expressed in **Eq. 1**.

### Thermal unfolding studied by circular dichroism (CD) and fluorescence spectroscopy

The thermal stability of the secondary structure of two porin variants, WT *Vh*ChiP and the W136A mutant, was examined using far-UV CD spectroscopy. The secondary structural elements of *Vh*ChiP were assessed from protein CD scans, with data collected from 190 to 260 nm. As shown in **Fig. 5A**, the CD spectra of *Vh*ChiP WT and the W136A mutant are essentially identical, showing a positive band at 198 nm and a negative band at 218 nm. Such characteristic bands are assigned to *β*-pleated sheets [26,27]. When *Vh*ChiP WT was subjected to heat denaturation (100 °C for 10 min), the band at 198 nm was shifted toward a more negative value, while the band at 218 nm was shifted upward, similar to the CD spectrum of the denatured protein reported previously [28]. The thermal unfolding profiles of ligand-free WT *Vh*ChiP and the W136A mutant, compared with those of the sugar-bound proteins, are shown in **Fig. 5B-C**, respectively. Fitting of the data using a Boltzmann sigmoidal function yielded the folding/unfolding curves and the midpoint temperature on each curve was denoted *T*_m_. All the fitted curves demonstrated an overall increment in the intensity when the temperature was progressively raised from 25 to 100 °C. The *T*_m_ values for the *Vh*ChiP variants in the absence and presence of ligands GlcNAc, chitobiose, -triose, -tetraose, -pentaose and -hexoase and maltohexaose (as a negative control) are summarized in **Table 1**. The representative CD profiles of *Vh*ChiP without and with chitohexaose, chitotetraose, GlcNAc and maltohexaose are presented in **Fig. 5A-B**, simplified for clarity. Our observations were that the addition of all chitooligosaccharides elevated the *T*_m_ of both WT *Vh*ChiP and the W136 mutant. The *T*_m_ of ligand-free W136A (59.3 °C) was 13.4 °C lower than that of the ligand-free WT channel (72.7 °C). Notably, the *T*_m_ of the sugar-protein complexes increased with increasing chain-length of chitooligosaccharide. Chitohexaose binding gave the highest *T*_m_ values, of 79.7 and 63.4 °C, for WT and mutant W136A respectively. The *T*_m_ was lowest with GlcNAc (74.1 °C for WT and 59.8 °C for the mutant). The difference in *T*_m_ between ligand-protein complex and the protein without ligand was calculated as Δ*T*_m_ (**Table 1**). In general, the Δ*T*_m_ values with all chitooligosaccharides were found to be significantly higher for WT than for the W136A mutant. The largest Δ*T*_m_ values were observed with the chitohexaose-WT and chitohexaose-W136A complexes, compared to their respective complexes with shorter chitooligosaccharides. With each ligand, the Δ*T*_m_ of W136A was significantly lower than that of WT.

**Fig. 5.**
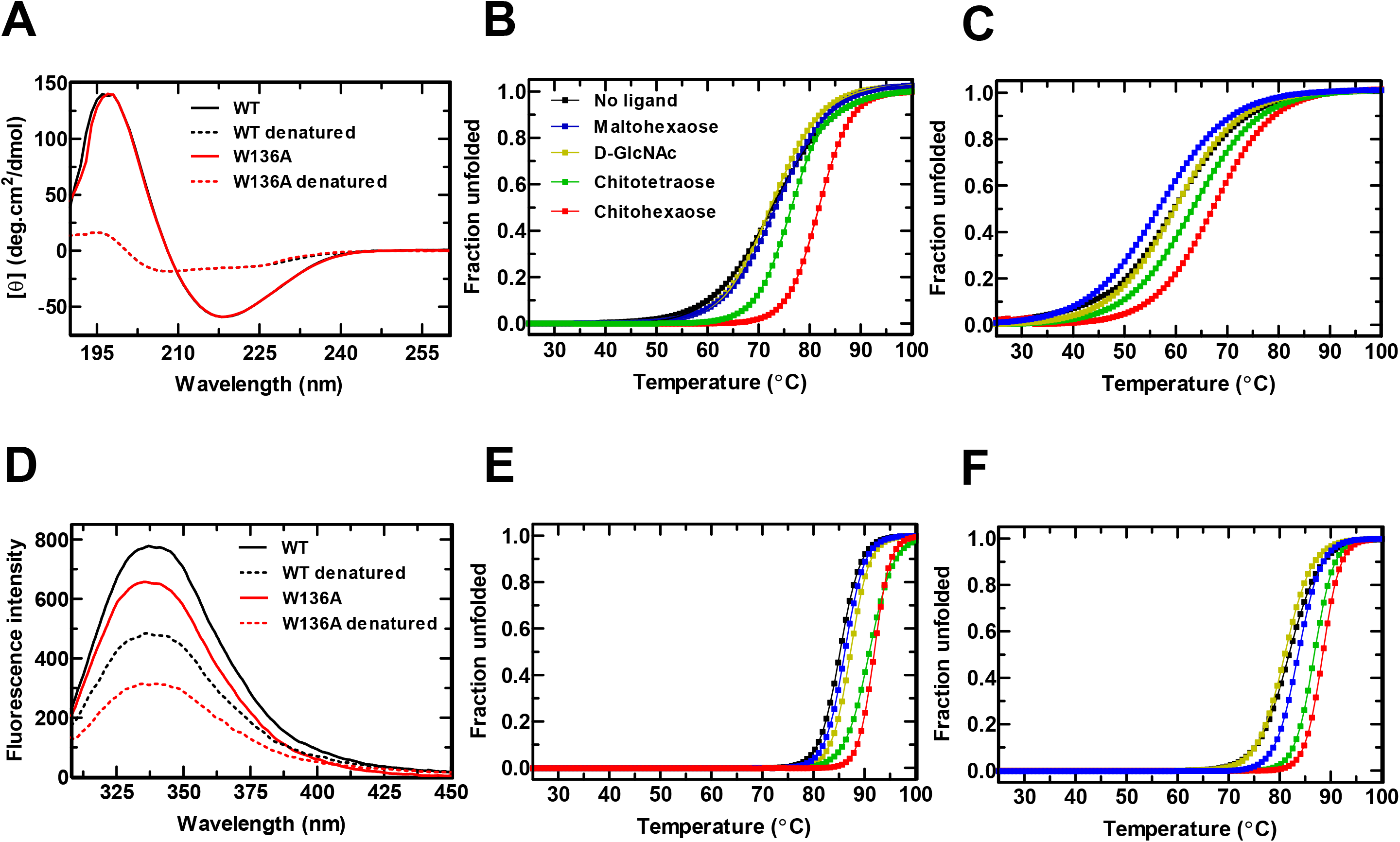
Thermal unfolding profiles of *Vh*ChiP as determined by CD (**A-C**) and fluorescence spectroscopy (**D-F**). (**A)** representative far-UV CD spectra of *Vh*ChiP acquired at 190-260 nm. Normalised thermal unfolding curves with and without sugar are shown in **(B)** for WT and (**C)** for W136A. Degrees of unfolding were calculated from CD intensity at 218 nm, the wavelength of greatest difference. WT or W136A (1 μM) was added with sugars (100 μM). (**D**) Representative Trp fluorescence spectra of *Vh*ChiP acquired using excitation at 295 nm and emission scanned from 300 to 450 nm. The normalised thermal unfolding curves of WT **(E)** and W136A **(F)** were obtained from the fluorescence spectra of the proteins (10 μM) in the absence or presence of sugars (1 mM). Maltohexaose was used as a control. For (**A**) and (**D**), the denatured proteins were obtained by boiling at 100 °C for 15 min before being subjected to CD or fluorescence scans using progressively increasing temperatures from 25 to 100 °C.

**Table 1.** Thermal unfolding analysis of *Vh*ChiP binding to chitooligosaccharides obtained from far-UV CD and fluorescence measurements. Each oligosaccharide (100 μM) was added to *Vh*ChiP WT and W136A, and spectral data acquired on gradual elevation of the temperature from 25 to 100 °C. The midpoint temperature (*T*_m_) values were obtained from plots of the fraction unfolded *vs*. temperature (°C). Δ*T*_m_ is the difference between the *T*_m_ values with and without ligand present. The values presented are mean ± SD obtained from three separate experiments. The slopes of the curve fits were obtained from the Boltzmann sigmoidal function available in GraphPad Prism *v.* 6.0.

| Ligand | Circular Dichroism |  |  |  |  |  | Fluorescence |  |  |  |  |  |
| --- | --- | --- | --- | --- | --- | --- | --- | --- | --- | --- | --- | --- |
|  | WT |  |  | W136A |  |  | WT |  |  | W136A |  |  |
| | $T_m$ ($^{\circ}$C) | $\Delta T_m$ ($^{\circ}$C) | *Slope* ($^{\circ}$C$^{-1}$) | $T_m$ ($^{\circ}$C) | $\Delta T_m$ ($^{\circ}$C) | *Slope* ($^{\circ}$C$^{-1}$) | $T_m$ ($^{\circ}$C) | $\Delta T_m$ ($^{\circ}$C) | *Slope* ($^{\circ}$C$^{-1}$) | $T_m$ ($^{\circ}$C) | $\Delta T_m$ ($^{\circ}$C) | *Slope* ($^{\circ}$C$^{-1}$) |
| No ligand | 72.7 ± 0.2 | 0.0 | 14.9 | 59.3 ± 0.4 | 0.0 | 8.1 | 85 ± 0.2 | 0.0 | 35.9 | 83.2 ± 0.2 | 0.0 | 26.4 |
| Chitohexaose | 79.7 ± 0.6 | +7.0 | 26.4 | 63.4 ± 0.2 | +4.1 | 11.2 | 93 ± 0.3 | +8.0 | 61.6 | 88 ± 0.3 | +4.8 | 51.5 |
| Chitopentaose | 78.4 ± 0.4 | +5.7 | 26.2 | 62.4 ± 0.1 | +3.1 | 9.8 | 91 ± 0.7 | +6.0 | 60.0 | 87 ± 0.2 | +3.8 | 49.1 |
| Chitotetraose | 76.5 ± 0.3 | +3.8 | 22.2 | 61.7 ± 0.6 | +2.4 | 9.6 | 90 ± 0.5 | +5.0 | 57.3 | 86 ± 0.4 | +2.8 | 48.6 |
| Chitotriose | 75.9 ± 0.2 | +3.2 | 21.9 | 60.1 ± 0.5 | +0.8 | 9.4 | 89 ± 0.1 | +4.0 | 56.1 | 85 ± 0.4 | +1.8 | 45.8 |
| Chitobiose | 74.8 ± 0.7 | +2.1 | 16.3 | 59.9 ± 0.3 | +0.6 | 9.4 | 89 ± 0.7 | +4.0 | 47.1 | 84 ± 0.3 | +0.8 | 43.6 |
| D-GlcNAc | 74.1 ± 0.3 | +1.4 | 13.8 | 59.8 ± 0.2 | +0.5 | 9.2 | 87 ± 0.5 | +2.0 | 44.7 | 82 ± 0.2 | -1.2 | 37.5 |
| Maltohexaose | 73.7 ± 0.6 | +1.0 | 14.0 | 56.8 ± 0.1 | -2.5 | 8.0 | 85 ± 0.7 | 0.0 | 40.1 | 82 ± 0.1 | -1.2 | 30.0 |

The temperature-induced unfolding of *Vh*ChiP WT and W136A was also carried out and monitored through intrinsic tryptophan fluorescence (**Fig. 5D-F**). The shifts of the fluorescence spectra on conversion of *Vh*ChiP from the folded to the unfolded state were monitored in the visible range of emission (wavelength 310 to 450 nm), with excitation at 295 nm (**Fig. 5D**). W136A showed a large reduction in fluorescence intensity, compared to the WT channel, and the denatured mutant showed a further reduction in the fluorescence intensity. To monitor the thermal stability of the tertiary structure of trimeric *Vh*ChiP, the fluorescence intensity in the same scan range was observed with gradual temperature increments from 65 to 100 °C. The normalised fractional unfolding curves of *Vh*ChiP WT (**Fig. 5E**), and W136A (**Fig. 5F**) in the presence of different chitooligosaccharides and maltohexaose were fitted by a Boltzmann sigmoidal function as specified in **Materials and Methods**, yielding the unfolding curves and allowing estimation of *T*_m_ values from the midpoints of the curves. Generally, a decrease in fluorescence intensity was observed when the protein was heated to denaturation. The results of thermal unfolding experiments obtained from the fluorescence measurements showed a trend similar to that observed in the CD experiments. *T*_m_ of ligand-free WT *Vh*ChiP was 85.0 °C, while that of the W136A mutant was 83.2 °C. As was observed from CD spectroscopy, both *T*_m_ and Δ*T*_m_ were elevated with increasing chain-length of chitooligosaccharide, the highest values being for the chitohexaose-protein complexes. The *T*_m_ and Δ*T*_m_ values of the ligand-bound WT protein were generally greater than those of the ligand-bound mutant.

We further analysed the slopes of the Boltzmann sigmoidal curve fits, which reflect cooperativity in the thermal unfolding processes. In the CD measurements, the ligand-free WT has a slope of 14.9 (**Table 1)**, while the slope is larger for the sugar-bound proteins, increasing with greater chain length of the chitooligosaccharides. The slope of the thermal unfolding curve for the unliganded W136A (slope = 9.8) is 1.5-fold lower than the value for the unliganded WT channel, and we found no substantial elevation in the value when different sugars were bound to the mutant channel.

For fluorescence measurements, the slope for the unbound WT channel is 35.9, while the slope for the unbound mutant was decreased 1.4-fold, to 26.4. Sugar binding was found to enhance the cooperativity of thermal unfolding transitions in both WT and W136A mutant, the slopes of the unfolding curves of the proteins bound to all chitooligosaccharides being considerably larger than those for the channels without bound ligands. The slopes consistently increase with increasing length of sugar chain, those with bound chitohexaose being greatest (61.6 for WT and 57.5 for W136A).

Small sugars of low affinity for *Vh*ChiP (GlcNAc) and the non-binding oligosaccharide maltohexaose did not alter the slopes for WT and W136A in the CD plots and slightly increased the values in the fluorescence plots.

### Binding studies using intrinsic tryptophan fluorescence spectroscopy

Intrinsic fluorescence changes were used to assess the binding affinity for various chitooligosaccharides of *Vh*ChiP and its mutant, (**Fig. 6**). When excited at 295 nm, ligand-free WT *Vh*ChiP and W136A emitted maximally (*F*max) at 350 nm. The magnitude of *F*max for the WT channel **(Fig. 5A**, 0 μM) was significantly higher than that for the W136A mutant (**Fig. 6B**, 0 μM). Titration of *Vh*ChiP WT and W136A mutant with chitohexaose (**Fig. 6A** **and** **6B**) enhanced the fluorescence emission in a concentration-dependent manner. Plots of relative changes in fluorescence intensity as a function of sugar concentration yielded saturation curves that could be fitted well by a one-site binding model (**Fig. 6C**). The fitted curves of the changes in fluorescence intensity (Δ*F*) *vs*. chitohexaose concentrations allowed the binding constant (*K*, M^−1^) to be calculated. The calculated binding energy (*ΔG*°_binding_) values are summarized in **Table 2**. Enhanced fluorescence signals were also observed when *Vh*ChiP WT (**Fig. 6D**) and W136A mutant (**Fig. 6E**) were titrated with chitotetraose, with the fitted curves shown in **Fig. 6F**. Clearly, W136A showed significantly lower *K* values for all chitooligosaccharides than those of the WT channel, indicating a lowering of the binding affinity by the Trp mutation. The equilibrium binding constant (*K*) obtained from fluorescence titration for the best substrate, chitohexaose, was reduced eight-fold (from 13 × 10^5^ M^−1^ in the WT channel to 1.7 × 10^5^ M^−1^ in the W136A mutant, **Table 2**). The *K* values for both WT and W136A shown in **Table 2** are in the order: chitohexaose > chitopentaose > chitotetraose > chitotriose > chitobiose > GlcNAc. When the WT (**Fig. 6G**) and W136A mutant (**Fig. 6H**) were titrated with an unrelated oligosaccharide (maltohexaose), there was no significant change in the fluorescence intensity, only scattered plots of Δ*F vs*. maltohexaose concentration (**Fig. 6I**), from which *K* values could not be obtained.

**Fig. 6.**
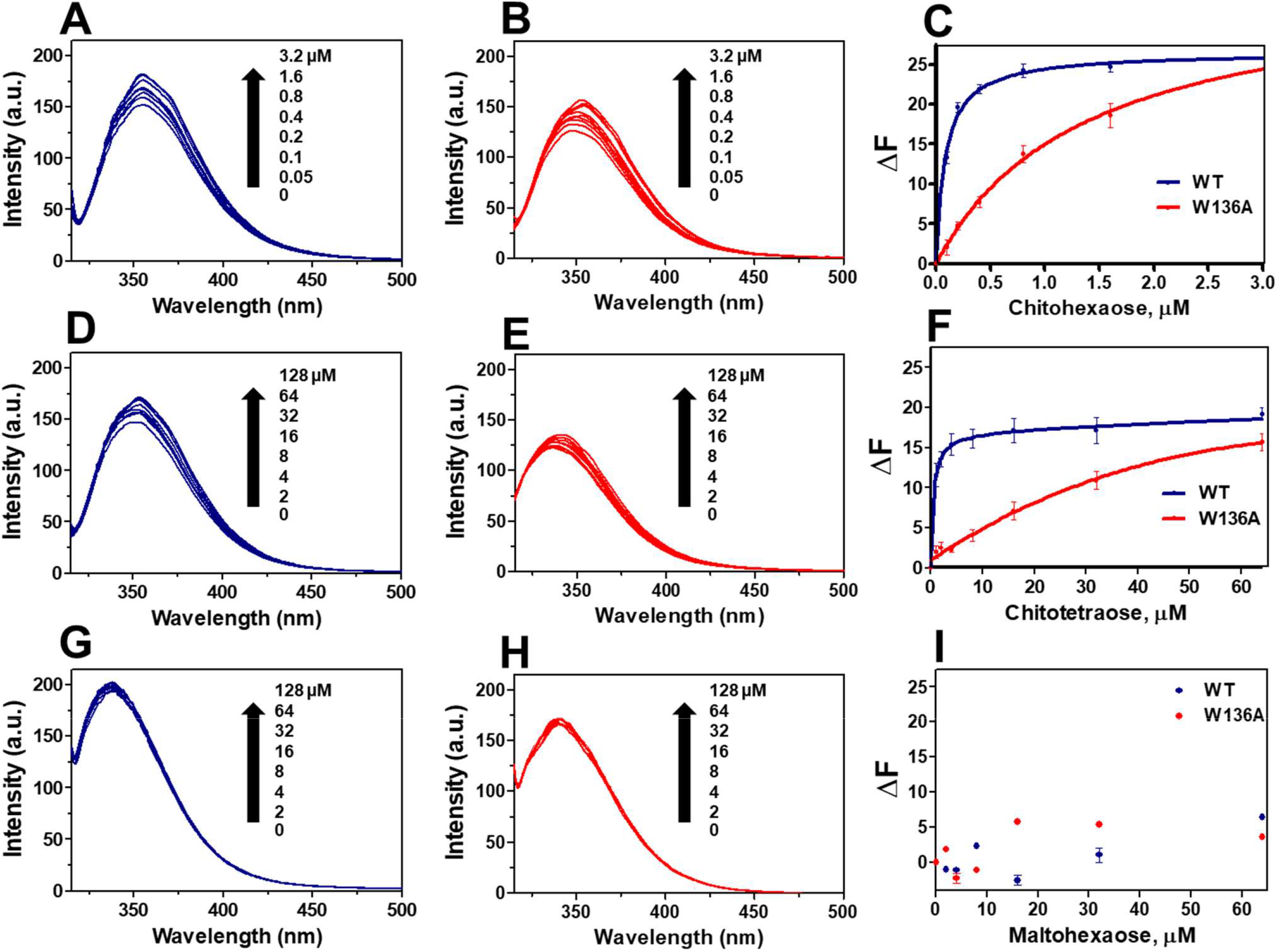
Tryptophan fluorescence titration of WT (blue) and W136A (red). (**A**) WT titrated with chitohexaose, (**B**) W136A with chitohexaose, and (**C**) the binding curves of WT and W136A with chitohexaose. (**D**) WT titrated with chitotetraose, **(E)** W136A titrate with chitotetraose, **(F**) the binding curves of WT and W136A with chitotetraose. (**G**) WT titrated with maltohexaose, (**H**) W136A titrated with chitotetraose, (**I**) the binding curves of WT and W136A with maltohexaose. The binding constants (*K*) were estimated from Michaelis-Menten plots of the fluorescence emission changes (Δ*F) vs*. sugar concentrations, according to **Eq. 4**.

**Table 2.** Determination of the binding affinity from intrinsic tryptophan fluorescence spectroscopy. *Vh*ChiP WT and W136A mutant were titrated with chitooligosaccharides of various chain lengths (*n* = 2 – 6). Values of Δ*G*°_binding_ were calculated from **Eq. 5**, while the binding constants (*K*, M^−1^) were obtained from the resultant binding curves, as defined by **Eq. 6**. The values presented are means ± SD obtained from at least three separate experiments.

| Ligand | WT |  | W136A |  |
| --- | --- | --- | --- | --- |
| | $K$<br>($\times 10^5 \text{ M}^{-1}$) | $\Delta G^{\circ}_{\text{binding}}$<br>($\text{kcal.mol}^{-1}$) | $K$<br>($\times 10^5 \text{ M}^{-1}$) | $\Delta G^{\circ}_{\text{binding}}$<br>($\text{kcal.mol}^{-1}$) |
| Chitohexaose | $13 \pm 0.5$ | $-8.3 \pm 0.2$ | $1.7 \pm 0.07$ | $-7.1 \pm 0.2$ |
| Chitopentaose | $3.8 \pm 0.2$ | $-7.6 \pm 0.2$ | $0.8 \pm 0.04$ | $-6.7 \pm 0.3$ |
| Chitotetraose | $1.4 \pm 0.01$ | $-7.0 \pm 0.1$ | $0.1 \pm 0.02$ | $-5.5 \pm 0.3$ |
| Chitotriose | $0.5 \pm 0.03$ | $-6.4 \pm 0.3$ | $0.02 \pm 0.01$ | $-4.5 \pm 0.1$ |
| Chitobiose | $0.1 \pm 0.04$ | $-5.5 \pm 0.2$ | $0.01 \pm 0.00$ | $-4.0 \pm 0.1$ |

Further temperature-dependence experiments were carried out using Trp fluorescence spectroscopy. The fluorescence spectra of WT *Vh*ChiP on titration with chitohexaose (**Fig. 7A)** and chitotetraose (**Fig. 7B)** were obtained at 25, 30, 35, 40, and 45 °C and their binding curves were evaluated using non-linear regression, allowing estimation of the dissociation constant (*K*_d_) and the binding constant (*K* = 1/*K*_d_). Secondary plots of log*K* against 1/*T* (temperature, Kelvin) for each binding curve were linear, yielding the enthalpy change (Δ*H*°) and the entropy change (Δ*S*°) according to the van’t Hoff equation (**Eq.6**). The Gibbs free energy can then be calculated from the relationship Δ*G*° = Δ*H*° − *T*Δ*S*° (**Eq. 7**). A plot of ln*K vs.* 1/*T* for WT and chitohexaose is shown in **Fig. 7C**, and for W136A and chitohexaose in **Fig. 7D**. The slope of these linear plots gives the term – Δ*H*°/*R* while the y-intercept equals Δ*S*°/*R*. The negative enthalpy changes (Δ*H*° = −38.3 ± 2.7 and −17.3 ± 1.0 kcal.mol^−1^, respectively: see inserts in **Figs. 7C** and **7D**) suggest that the binding of chitohexaose to the *Vh*ChiP variants is an exothermic process and involves hydrogen-bond interactions. The significantly greater enthalpy change on binding implies that the complex of chitohexaose with WT is more stable than that with W136A. The entropy change (Δ*S*°) expresses the change in the degree of randomness of the system. The degree of disorder, indicated by the term −TΔ*S*°, was calculated for WT *Vh*ChiP as +28.8 ± 1.3 kcal.mol^−1^ and for the W136A mutant as +9.5 ± 1.0 kcal.mol^−1^. This large positive −TΔ*S*° term suggests that the binding of chitohexaose to the protein has a negative Δ*S*^°^ at 25 °C, corresponding to an increase in the structural order of the protein upon sugar binding.

**Fig. 7.**
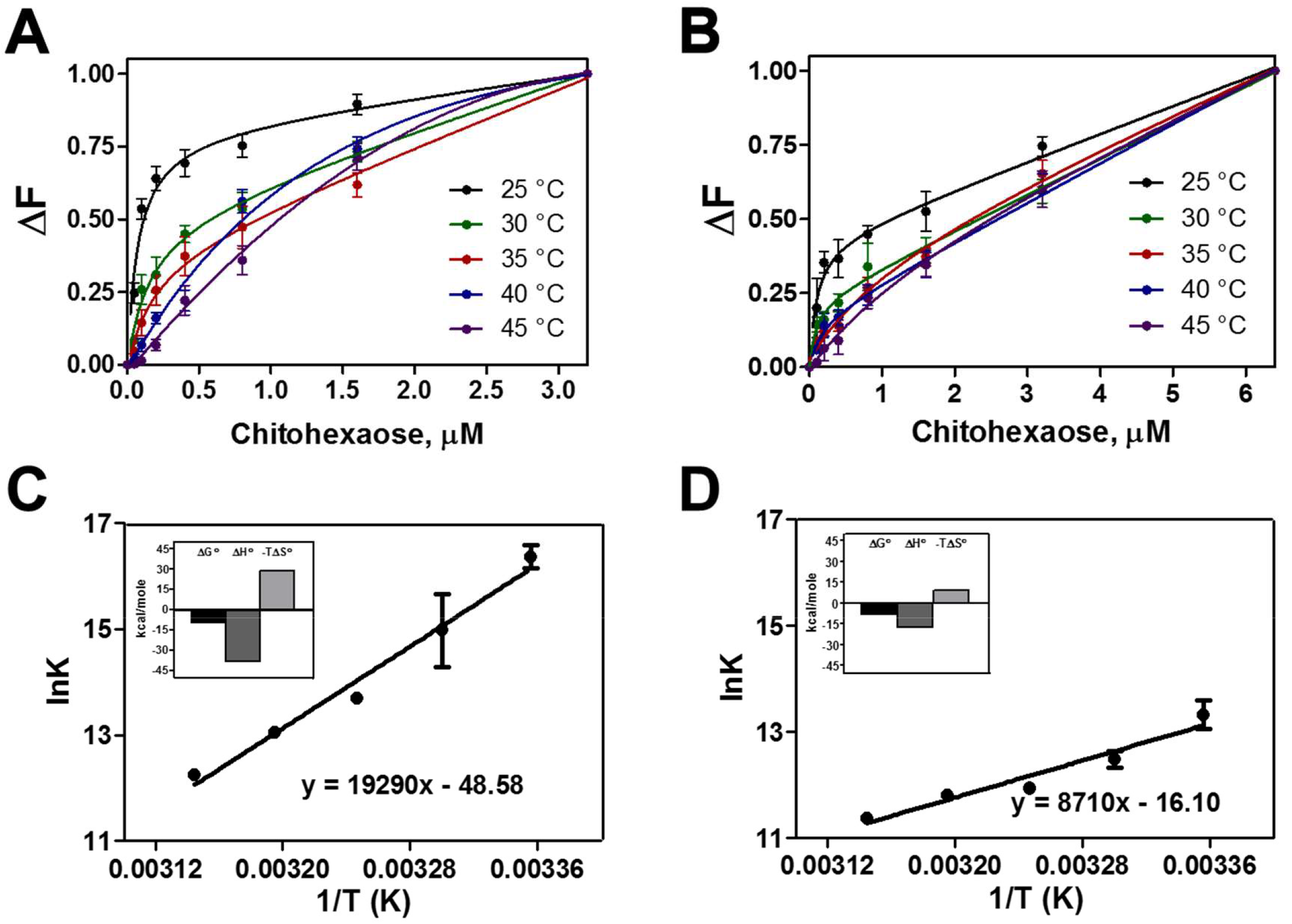
Thermodynamics of interactions of WT and the W136A mutant with chitohexaose, studied by intrinsic fluorescence titration measurements. The plots for WT (A) and W136A (B) are presented separately. Titrations were conducted at 25, 30, 35, 40 and 45 °C. Binding constants (*K*) were estimated from plots of Δ*F vs*. chitohexaose concentration. Panels **C** and **D** show van’t Hoff plots, of the logarithm of the binding constants versus the reciprocal of temperatures at which the titrations were performed, using data from panels A and B, respectively. The inset panels show comparisons of free energy change (Δ*G*°_binding_), enthalpy change (Δ*H*°) and the entropy change (−*T*Δ*S*°) for binding of chitohexaose by each protein.

### Binding studies using single-channel electrophysiology

Single-channel recordings were made to study chitooligosaccharide interactions with the *Vh*ChiP channel (both WT and W136A mutant) at the single-molecule level. **Figure 8A** shows that chitohexaose interacted strongly with WT *Vh*ChiP, causing the occlusion of ion flow in a concentration-dependent manner. The histogram analysis showed a gradual increase in occlusions, from single subunits to two or three subunits, as the sugar concentration was increased from 1.0 to 5.0 μM **(Fig. 8A**, **inset).** Chitotetraose could also block the channel (**Fig. 8B**), but blocking events were short-lived, like those observed with the W136A mutant blocked by chitohexaose, as shown in **Fig. 8C**. On addition of chitotetraose to the W136A mutant, blocking events were rarely observed (**Fig. 8D**). Histogram analysis showed only a slight reduction in the monomeric conductance of *Vh*ChiP with added chitotetraose and of the W136A mutant with added chitohexaose (**Fig. 8B** and **8C**, **inset)**. No change in monomeric subunit conductance was detectable in the W136A mutant on addition of chitotetraose **(Fig. 8D**, **inset)**. **Table 3** summarizes the kinetic analysis obtained from the sugar-dependent blocking events. These parameters include the equilibrium binding constant (*K*) and the on-rate (*k*_on_) and off-rate (*k*_off_) constants. Chitohexaose saturated the channel at concentrations above 2.5 μM, and *k*_on_ for chitohexaose was estimated from the number of blocking events obtained over a low concentration range (0 − 2.5 μM) to avoid overlapping of the monomeric, dimeric and trimeric blockade.

**Fig. 8.**
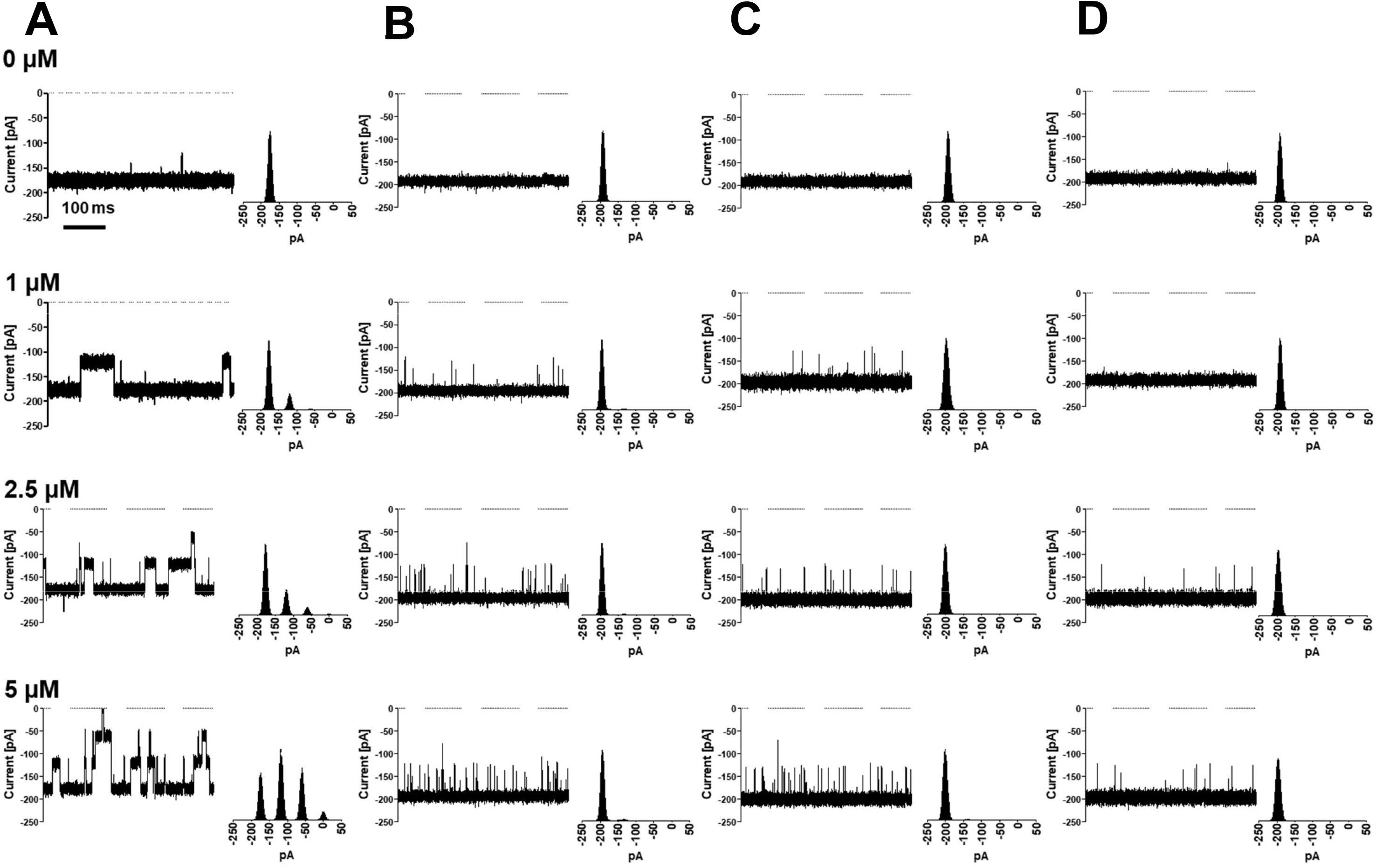
Time-resolved single-channel recordings of sugar-channel interactions. A single molecule of *Vh*ChiP was reconstituted in an artificial phospholipid membrane, then titrated with chitooligosaccharide. **A**, WT titrated with chitohexaose; **B**, WT titrated with chitotetraose; **C**, W136A titrated with chitohexaose; **D**, W136A titrated with chitotetraose. Ion current traces were acquired at −100 mV with the sugar added on the *cis* side. Insets are histogram analyses of each corresponding ion trace, reflecting sugar-induced closure of the protein channel.

**Table 3.**
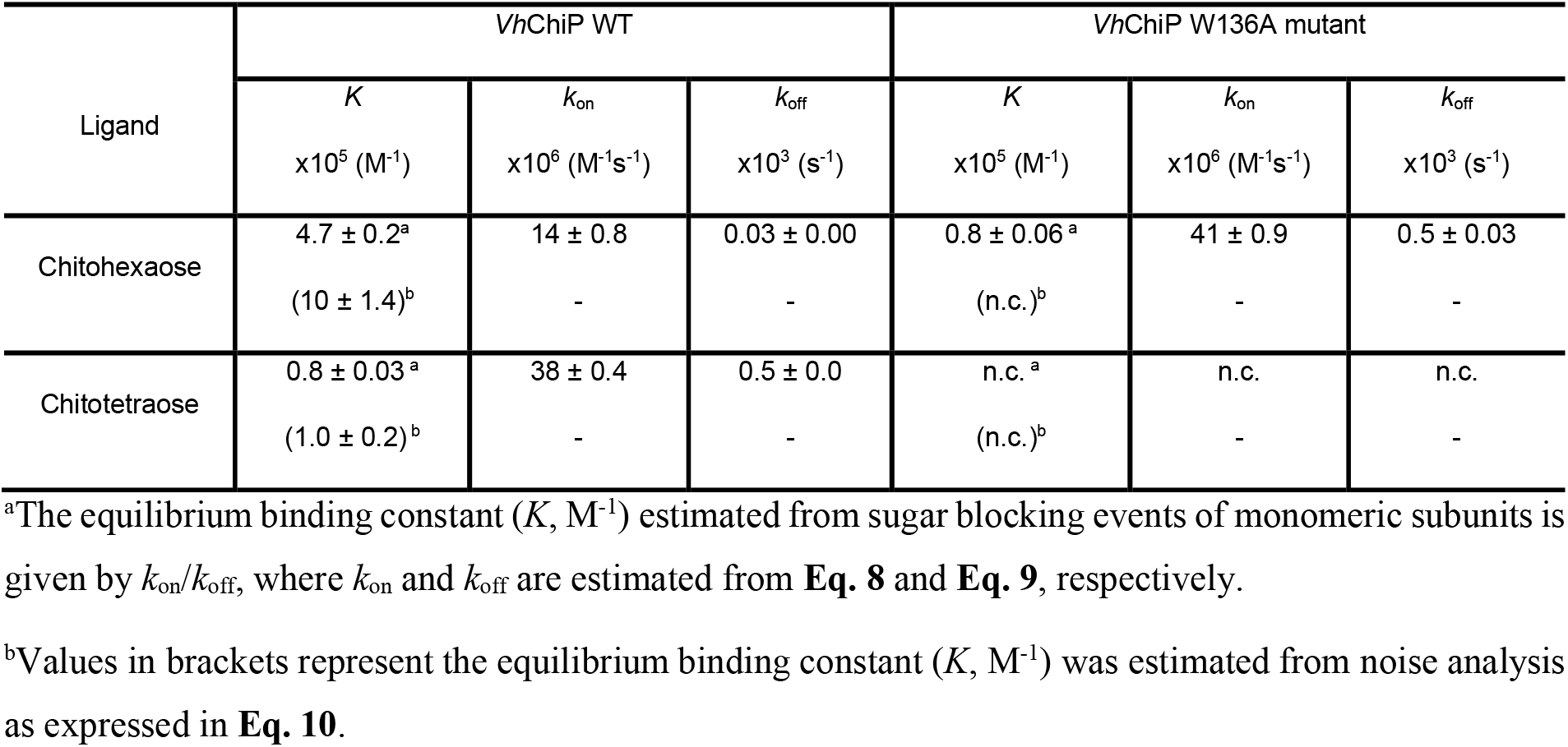
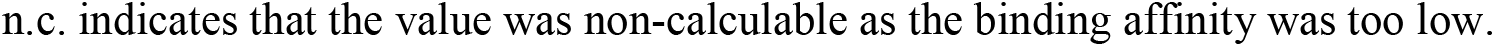
The kinetic parameters for chitooligosaccharide binding to *Vh*ChiP WT and W136A, obtained from single-channel recordings. The values presented are means ± SD obtained from three separate measurements.

| Ligand | *VhChiP* WT |  |  | *VhChiP* W136A mutant |  |  |
| --- | --- | --- | --- | --- | --- | --- |
| | $K$<br>$\times 10^5 (\text{M}^{-1})$ | $k_{\text{on}}$<br>$\times 10^6 (\text{M}^{-1}\text{s}^{-1})$ | $k_{\text{off}}$<br>$\times 10^3 (\text{s}^{-1})$ | $K$<br>$\times 10^5 (\text{M}^{-1})$ | $k_{\text{on}}$<br>$\times 10^6 (\text{M}^{-1}\text{s}^{-1})$ | $k_{\text{off}}$<br>$\times 10^3 (\text{s}^{-1})$ |
| Chitohexaose | $4.7 \pm 0.2^{\text{a}}$ | $14 \pm 0.8$ | $0.03 \pm 0.00$ | $0.8 \pm 0.06^{\text{a}}$ | $41 \pm 0.9$ | $0.5 \pm 0.03$ |
| | $(10 \pm 1.4)^{\text{b}}$ | - | - | (n.c.)<sup>b</sup> | - | - |
| Chitotetraose | $0.8 \pm 0.03^{\text{a}}$ | $38 \pm 0.4$ | $0.5 \pm 0.0$ | n.c. <sup>a</sup> | n.c. | n.c. |
| | $(1.0 \pm 0.2)^{\text{b}}$ | - | - | (n.c.)<sup>b</sup> | - | - |

<sup>a</sup>The equilibrium binding constant ( $K$ , $\text{M}^{-1}$ ) estimated from sugar blocking events of monomeric subunits is given by $k_{\text{on}}/k_{\text{off}}$ , where $k_{\text{on}}$ and $k_{\text{off}}$ are estimated from **Eq. 8** and **Eq. 9**, respectively.
<sup>b</sup>Values in brackets represent the equilibrium binding constant ( $K$ , $\text{M}^{-1}$ ) was estimated from noise analysis as expressed in **Eq. 10**.
n.c. indicates that the value was non-calculable as the binding affinity was too low.

As shown in **Table 3**, the *K* value obtained for WT *Vh*ChiP titrated with chitohexaose (4.7 ×10^−6^ M^−1^) was six-fold greater than that for W136A titrated with the same substrate (0.8 × 10^−6^ M^−1^). It is noteworthy that the binding constants reported from single channel recordings were two-to three-fold lower than the values obtained from fluorescence titration. Values of *K* estimated from *k*_on_/*k*_off_ were expected to be lower than the actual values because of a limitation of our electrophysiology setup, in which short-lived blocking events, with a residence time of <100 μs, were essentially invisible. In addition, such events are beyond the detection limit of the data-analysis software (Clampfit *v*. 10.0), with rare overlapping monomeric and dimeric blocking leading to underestimation of the unresolvable blocking events. Nevertheless, the *K* values for titrations of WT with chitohexaose (*K* = 10 ± 1.4 × 10^6^ M^−1^) and chitotetraose (*K* = 1.0 ± 0.2 ×10^6^ M^−1^) (**Table 3**, **values in brackets**) obtained from noise analysis (**Eq. 10**) were comparable to the values obtained from fluorescence enhancement.

## Discussion

The substrate-specific porins of Gram-negative bacteria are responsible for the transport of small molecules such as sugars, amino acids and peptides, which are required for the bacteria as their carbon and nitrogen sources. Oligomeric substrates usually interact with affinity sites that may contain multiple binding sites, depending on the natural architecture of their substrates. Maltoporin (LamB) is among the best-studied examples of substrate-specific porins that contain multiple binding sites for long-chain maltooligosaccharides [29,30], other examples being ScrY for sucrose [31], CymA for cyclodextrin [32], OprB for monosaccharides [33] and OprD for amino acids and peptides [34]. We recent studied the chitooligosaccharide-specific porin from *V. campbellii (Vh*ChiP). The protein, expressed in *E. coli*, was shown to form stable trimeric channels in artificial phospholipid membranes, with an average conductance of 1.8 ± 0.3 nS [11,12] and to exhibit the greatest affinity for chitohexaose [23,18]. It was demonstrated by the structures of *Vh*ChiP-substrate complexes that the surface of the *Vh*ChiP channel lumen contains multiple GlcNAc-binding sites that can fully accommodate chitohexaose, while chitotetraose binds with lower affinity in the central area of the protein channel (**Fig. 2B**, **the sugar backbone in black**). Notably, observable scattered electron density at the end sites +1 and +6 may indicate the movement of the chitotetraose chain along the sugar-binding sites to achieve translocation through the protein channel. We previously reported the use of statistical analysis of stochastic fluctuations of ion current through the *Vh*ChiP channel reconstituted in bilayer lipid membranes, which indicated that *Vh*ChiP contains multiple binding sites and that sugar-channel and/or sugar-sugar interactions helped to direct the movement of the sugar chain through the channel [35,36].

In this study we employed both CD and fluorescence spectroscopy to demonstrate that the ligand-bound state of *Vh*ChiP had a higher *T*_m_ than the ligand-free state. The results clearly indicated the protein channel was more resistant to thermal denaturation when bound to its sugar substrates, the degree of heat tolerance being greater with increasing binding affinity. Such observations have been reported in previous studies: for example, it was shown by Fukamizo’s group that chitosan hexamer helped to stabilize the structure of the glucosamine-recognizing β3 conglycinin from soybean [37].

*Vh*ChiP mutant W136A showed significantly reduced *T*_m_ values in both the unliganded and ligand-bound states (**Table 1**), as well as decreased binding affinity for all chitooligosaccharides. The considerable reduction in *T*_m_ values with the Ala substitution of Trp^136^ observed in the CD and Trp thermal unfolding profiles, together with lower temperature required for subunit dissociation as seen on SDS-PAGE analysis, emphasized the important role of Trp^136^ in stabilizing the sugar-channel complex. In our structural dissection, the mutation abolished hydrophobic interactions between the sugar and the channel, thus destabilizing the overall structure of the protein. Numerous intermediate states are usually seen from thermal folded/unfolded curves when a protein contains multiple domains with diverse conformations [38]. However, the thermal unfolding profiles obtained from both methods shown in **Fig. 5B-C** (CD) and **Fig. 5E-F** (Trp fluorescence) indicate a two-state transition, with no discrete intermediate states between completely folded and completely unfolded *Vh*ChiP.

The observed slope of the fractional unfolding curves, larger than 1, suggests multiple conformational states with cooperativity during unfolding of the *Vh*ChiP channels. Strong cooperative effects in the unfolding transitions at the three-dimensional level are obviously indicated by the steep slopes in the fluorescence plots (**Fig. 5E-F**). In contrast, a weaker cooperative effect is seen from the shallower slopes of the CD plots (**Fig. 5B-C**), which indicates a lower degree of cooperativity during the unwinding of the secondary elements (mainly *β*-strands and connecting loops). Sugar binding clearly enhanced the cooperativity of protein unfolding, and the effects were amplified with increasing length of the chitooligosaccharide (**Table 1**). Reduced *T*_m_ and slope values in the thermal unfolding curves of the W136A mutant, in fractional unfolding curves from both CD and fluorescence, also emphasize the role of Trp^136^ in thermal stabilization and unfolding cooperativity. The effects of the mutation are more pronounced at the level of the tertiary structure than the secondary structure of *Vh*ChiP. Honda *et al* (39) and Fukamizo *et al* (40) reported similar phenomena in the effects of chitosan oligomers on thermostability and cooperativity of the unfolding transitions of a *Streptomyces sp.* N174 chitosanase. Although substrate binding showed good fitting by a single-site model, our previous statistical analysis of stochastic current fluctuations revealed that the rate of sugar trapping events inside the *Vh*ChiP channels was enhanced when the other neighbouring monomers had been pre-occupied by sugar molecules and such so-called memory effects had been suggested as a possible strategy for *Vh*ChiP to enhance sugar transport (36). In this study, we observed that the presence of sugar (especially chitohexaose) inside the *Vh*ChiP channels helped to protect the protein trimers against thermal denaturation through its cooperative effects.

For both WT and the W136A mutant the *K* values increased with increasing chain length of the bound chitooligosaccharide. A previous study of the effects of chitooligosaccharide binding to a single channel in artificial phospholipid membranes by analysis of time-resolved blockade events showed that chitooligosaccharides (GlcNAc)_*n*_ of greater chain length, such as those with *n* = 4, 5 or 6, have high specificity for trimeric *Vh*ChiP [17]. Substrate binding by *Vh*ChiP involves mainly hydrophobic interactions and hydrogen bonds, which are typical of observed sugar-channel interactions [33,41,42].

Chitooligosaccharide chain-length clearly affected the free energy change on binding (Δ*G*°_binding_). Overall, Δ*G*°_binding_ became increasingly negative as *n* increased from 1 to 6, following the trends of the *T*_m_ changes obtained from CD experiments. The value of the enthalpy term (−Δ*H*°) for the chitohexaose-WT interaction (−38.3 ± 2.7 kcal.mol^−1^) was two-fold greater than with W136A (−17.3 ± 2.1 kcal.mol^−1^) and the positive entropic term (−TΔ*S*°) for the chitohexaose-WT interaction (+28.8 ± 1.3 kcal mol^−1^) was three-fold higher than the value with W136A (+9.5 ± 1.0 kcal.mol^−1^). A previous study suggested that the positive value of −TΔ*S*° was related to hydrophobic interactions and conformational changes around the sugar-binding sites, while the negative value of Δ*H*° implies the involvement of hydrogen bonds [43]. The lower free energy of binding for this sugar was predominately the result of hydrophobic effects that reflected the loss of a hydrophobic contribution due to the removal of the aromatic side chain of Trp136, while H-bond interactions had minor effects on the binding energy.

Finally, our findings, which elucidated the effects of chitooligosaccharide binding on the stabilization of the structural elements of the chitin uptake channel, suggest the feasibility of exploiting chitooligosaccharides as starting materials for chemo/enzymatic synthesis of sugar-based analogs using a series of reactions involving chemical modifications and glycoside hydrolase/glycosyl transferase enzymes. The newly-synthesized sugars that form stable complexes with sugar uptake channels such as chitoporin will be examined and further optimized, with a view to using them as potential antimicrobial agents to combat *Vibrio* infections.

## Materials and Methods

### Structural analysis of chitooligosaccharide-binding sites

Details of the sugar-channel interactions were analysed using the crystal structures of WT *Vh*ChiP complexed with chitohexaose (pdb id 5MDR) and chitotetraose (pdb id 5MDS), obtained by X-ray crystallography as reported previously [24]. Hydrophobic and hydrogen bonds between the protein and the ligands were generated by LIGPLOT [44] within 3.5 Å distance. The 2*F*_o_-*F*_c_ maps of chitooligosaccharides inside the channel were generated by COOT with σ level of 2.0. The sugar-binding sites are designated affinity sites +1, +2, +3, +4, +5, +6, following the description reported for the crystal structures [24]. The structures were visualised and displayed by PyMol (education *v.* 1.3) (www.pymol.org).

### Expression and purification of *Vh*ChiP variants

The *chip* gene from *V. campbellii* was cloned as described in our previous study [17]. The W136A mutant was generated by a PCR-based strategy using the full-length *chip* gene in the pET23d(+)/*Vh*ChiP construct as DNA template, as described by Chumjan *et al*. [45]. *E. coli* host strain BL21 (DE3) Omp8 Rosetta was genetically engineered to carry defective genes encoding major outer membrane porins OmpA, OmpC, OmpF and LamB, making it suitable for production of an exogenous porin [46, 47].

Expression of wild-type *Vh*ChiP and the W136A mutant was carried out as described previously [17]. Recombinant WT and the W136A mutant were expressed and purified, following the protocol originally described by Garavito and Rosenbusch [48]. In brief, the *E. coli* strain BL21 (Omp8) Rosetta cells, transformed with the recombinant plasmid pET23d (+) carrying the WT or mutated *chip* gene, were grown at 37 °C in Luria-Bertani (LB) liquid medium containing 100 μg mL^−1^ ampicillin, 25 μg mL^−1^ kanamycin and 1% (w/v) glucose. At an *OD*_600_ of 0.6 - 0.8, isopropyl *β*-D-thiogalactoside (IPTG) was added to a final concentration of 0.5 mM. Cell growth was continued for a further 6 h and cells were then harvested by centrifugation at 4,500 × g at 4 °C for 20 min. The cell pellet was resuspended in 20 mM Tris-HCl, pH 8.0, containing 2.5 mM MgCl_2_, 0.1 mM CaCl_2_, 10 μg mL^−1^ DNase and 10 μg mL^−1^ RNase A. Cells were lysed by sonication on ice for 10 min (30% duty cycle; amplitude setting 20%) using a Sonopuls Ultrasonic homogenizer with a 6-mm diameter probe. *Vh*ChiP was extracted with sodium dodecyl sulphate (SDS), based on the method of Lugtenberg and Alphen [49]. Briefly, 20% (w/v) SDS stock solution was added to the lysed cell suspension to obtain a 2% (w/v) final concentration, followed by incubation at 50 °C for 1 h with gentle stirring and centrifugation at 40,000 × g at 4 °C for 60 min. WT and mutant *Vh*ChiPs were extracted from the pellets, which were enriched in outer membranes, in two steps. In a pre-extraction step, the pellet was washed with 15 mL of 0.125% (v/v) *n*-octyl POE (*n*-octylpolyoxyethylene) in 20 mM sodium phosphate buffer pH 7.4 (ALEXIS Biochemicals, Lausen, Switzerland), homogenized with a Potter-Elvehjem homogenizer, incubated at 37 °C for 60 min then centrifuged at 100,000 × g, 4 °C for 40 min. In the second step, the pellet from centrifugation at 100,000 × g was resuspended in 10 - 15 mL of 3% (v/v) *n*-octyl POE in 20 mM sodium phosphate buffer pH 7.4, then homogenized with a Potter-Elvehjem homogenizer and incubated at 37 °C for 60 min, followed by further centrifugation at 100,000 × g, 4 °C for 40 min. The detergent was then exchanged with 0.2% (v/v) LDAO (*N,N*-lauryldimethylamine oxide, Sigma-Aldrich) by dialysis and the supernatant subjected to ion-exchange chromatography on a HiTrap Q HP prepacked column (5 × 1 mL), connected to an ÄKTA Prime plus FPLC system (GE Healthcare Life Sciences, Life Sciences Instruments, ITS (Thailand) Co., Ltd., Bangkok, Thailand). Bound proteins were eluted with a linear gradient of 0 - 1 M KCl in 20 mM sodium phosphate buffer, pH 7.4 containing 0.2% (v/v) LDAO. The purity of the eluted proteins was checked by SDS-PAGE and verified by immunoblotting. Fractions containing only *Vh*ChiP were pooled and the protein concentration was determined using the Pierce BCA protein assay kit (Bio-Active Co., Ltd., Bangkok, Thailand).

### Temperature-induced subunit dissociation

The reaction mixture (200 μL) containing 10 μM freshly purified *Vh*ChiP in 20 mM sodium phosphate buffer, pH 7.5, 0.1% (v/v) LDAO and either no addition or 100 μM oligosaccharide was pre-incubated on ice in an Eppendorf ThermoMixer® Comfort with gentle shaking for 1 h. 15-μL aliquots of the solution (18 μg protein) were added to 5 μL of sample loading buffer to give a final concentration of 2% (w/v) SDS and incubated at temperatures of 30, 40, 50, 60, 70, 80, 90 and 100 °C for 10 min. 9 μg of each protein complex was analysed by SDS-PAGE on 10% polyacrylamide gels, which were then stained with Coomassie Blue.

### Molecular weight determination of trimeric and monomeric *Vh*ChiP

Molecular weights of trimeric and monomeric of *Vh*ChiP were determined by gel filtration chromatography using a Superdex S-200 column (1.6 cm × 87 cm) connected to an ÄKTA™ Pure FPLC system (Bang Trading 1992 Co. Ltd., Bangkok, Thailand) with a flow rate of about 1.5 mL.min^−1^. Standard proteins of known molecular weight were dissolved in the equilibration buffer containing 20 mM sodium phosphate pH 7.4, 100 mM NaCl and 0.1% (w/v) LDAO. Blue dextran (1000 kDa; 0.8 mg) was used to obtain the void volume (*V*_o_), while DNP-lysine (348.7 Da; 0.35 mg) was used to calculate the bed volume of the stationary phase (*V*_t_), and the elution volume (*V*_e_) of each protein sample was characterized by the average distribution constant, (*K*_av_) expressed in **Eq. 1** [50]:

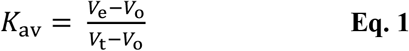

A calibration curve of native proteins was created by plotting *K*_av_ *vs.* logarithm of the corresponding molecular weight of the standard protein and was used to estimate the molecular weight of *Vh*ChiP incubated at 30 and 100 °C for 15 min. The standard proteins used were glucose oxidase (160 kDa; 4.0 mg applied), bovine serum albumin (66 kDa; 6.8 mg), ovalbumin (43 kDa; 6.6 mg) and lysozyme (14 kDa; 2.5 mg).

### Thermal unfolding observed by Circular Dichroism (CD)

Far-ultraviolet CD spectra were obtained with freshly purified *Vh*ChiP in 20 mM phosphate buffer, pH 7.5 and 0.2% (v/v) LDAO, using a Jasco J-815 Circular Dichroism Spectrometer (J & P Jasco Products, Co. Ltd., Bangkok, Thailand). Scans were carried out at 25 °C and the data acquired over the wavelength range 190 – 260 nM to determine the molar ellipticity [θ] profile of the secondary structural elements of the *Vh*ChiP WT and W136A mutant.

The expression defining molar ellipticity [θ] is given by **Eq. 2**:

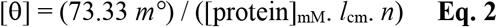

where [θ] is the molar ellipticity in deg. cm^2^. dmol^−1^, *n* is the number of amino acids in the polypeptide chain, *m*° is the measured ellipticity and *l*_cm_ is the path length in centimetres [**51,52]**. The intensity of standard CSA (non-hygroscopic ammonium (+)-10-camphorsulphonate) at wavelength 290 nm is approximately 45 units, giving the calculated conversion factor from 3300/CSA intensity using the above equation to be 73.33. From its corresponding nucleotide sequence, the number of amino acid residues per *Vh*ChiP monomer was predicted to be 352.

Molar ellipticity changes in the absence and presence of chitooligosaccharides were then monitored continuously at 220 nm while the solution temperature was raised from 25 to 100 °C at a rate of 1 °C min^−1^ by a temperature controller (PTC-423L, J & P Jasco Products, Co. Ltd.). Protein (1 μM) and *D*-GlcNAc and chitooligosaccharides (100 μM) were mixed in a 200-μL quartz cuvette. The fractional unfolding of the protein was calculated from the molar ellipticity [θ] difference between the fully folded and the fully unfolded states of the protein as expressed in **Eq. 3** (53).

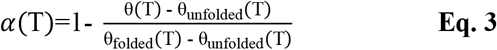

In **Eq.3**, α(T) is the fraction unfolded, θ(T) the measured ellipticity at a given temperature. θ_unfolded_(T) and θ_folded_(T) are the ellipticities of the unfolded and the folded state of the *Vh*ChiP trimers, respectively.

The fractional unfolded value was then plotted against temperature. The theoretical curves were fitted to the experimental data obtained by a non-linear least square fitting procedure. The *T*_m_ value for each unfolded state was obtained by direct fitting using the Boltzmann sigmoidal function available in GraphPad Prism *v.* 6.0 (GraphPad Software, California, USA).

### Thermal unfolding measurement by fluorescence spectroscopy

Fluorescence measurements were made with a Jasco FP-6200 Fluorescence spectrometer (J & P Jasco Products, Co. Ltd., Bangkok, Thailand), with *Vh*ChiP dissolved in 20 mM phosphate buffer, pH 7.5 and 0.2% (v/v) LDAO. Fluorescence scans were carried out with excitation at 295 nm and emission measurement at 300 – 450 nm. The fluorescence intensity was plotted against wavelength to obtain the fluorescence emission spectrum. For titration experiments, the emission wavelength that gave the maximum intensity (*F*_max_), 340 nm, was chosen and for each sugar titration, the solution temperature was raised from 25 to 100 °C at a rate of 1 °C min^−1^ by a temperature controller (PTC-423L, J & P Jasco Product, Co. Ltd.). The excitation and emission slit widths were 10 and 20 nm, respectively. Protein (10 μM) and *D*-GlcNAc and chitooligosaccharides [(GlcNAc)_n_, *n* = 2-6] (1 mM) were mixed in a 200-μL quartz cuvette. The fraction of unfolded protein was calculated from the fluorescence emission difference between folded and fully unfolded proteins. This fraction was then plotted against temperature. The theoretical curve was fitted to the experimental data by a non-linear least-square fitting procedure. *T*_m_ values for each unfolding process were obtained by direct fitting using the Boltzmann sigmoidal function available in GraphPad Prism *v.* 6.0.

### Binding studies using an intrinsic tryptophan fluorescence assay

Purified *Vh*ChiP (2.0 μM, in 20 mM phosphate buffer, pH 7.4 and 0.2% (v/v) LDAO was titrated with chitooligosaccharides (a concentration range of 0-3.2 μM for chitohexaose, 0-128 μM for chitotetraose and chitopentaose and 0-256 μM for chitobiose and chitotriose) at 25 ± 1 °C. Changes in the intrinsic tryptophan fluorescence intensity were monitored directly with a LS-50 fluorescence spectrometer (Perkin-Elmer Limited, Thailand). The excitation wavelength was set to 295 nm and emission spectra were collected over the range 300 – 450 nm, with excitation and emission slit widths of 5 and 10 nm, respectively. The spectrum of each protein was corrected for the blank. Binding curves were evaluated with a nonlinear regression function using a model based on a single binding-site, available in Prism *v*.6.0. To estimate the dissociation constant, relative fluorescence Δ*F* = *(F*_max_ − *F*_min_) was plotted as a function of sugar concentration, yielding a rectangular hyperbolic curve. This curve allows calculation of the dissociation constant for the chitooligosaccharides using a single-site binding model, according to **Eq. 4** [54, 55, 56]:

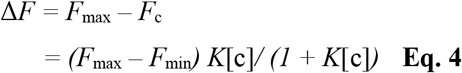

Δ*F* is the difference between fluorescence intensity before and after addition of the sugar ligand; *F*_max_ refers to the maximum emission intensity in the absence of sugar; *F*_min_ is the minimum emission intensity; *F*_c_ is the emission intensity at a given concentration of ligand; [c] is the concentration of ligand; and *K* is the equilibrium binding constant (M) [57]. The free energy change on binding (Δ*G*°_binding_) can be estimated from **Eq. 5**:

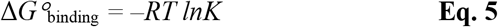

At equilibrium under conditions of constant pressure, the binding constant *K* is related to the standard Gibbs free-energy change (Δ*G*°_binding_) of the reaction, where *R* is the gas constant (1.9872 cal mol^−1^ K^−1^) and *T* is the temperature (in Kelvin).

### Determination of thermodynamic parameters of chitooligosaccharide binding using fluorescence spectroscopy

Titrations of chitohexaose into *Vh*ChiP WT and W136 mutant were carried out at temperatures of 25, 30, 35, 40 and 45 °C. Thermodynamic parameters were calculated to characterize the interactions between the *Vh*ChiP channels and chitohexaose. The thermodynamic parameters were determined under the standard condition and expressed by the van’t Hoff equation:

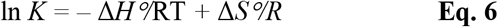

where Δ*S*° is the entropy change, *K* is the equilibrium binding constant at the corresponding temperature [58] and *R* is the gas constant (1.9872 cal mol^−1^ *K*^−1^). The enthalpy change (Δ*H*°) and the entropy change (Δ*S*°) were obtained from the slope of the van’t Hoff curve and y-intercept of the fitted curve of *ln K* against 1/*T*, respectively. The free energy change on binding (Δ*G*°_binding_) was estimated from **Eq. 7**:

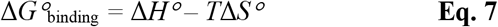

The van’t Hoff binding curves were plotted in Prism *v.* 6.0.

### Single molecule electrophysiology

Black lipid bilayer (BLM) measurements and single channel analysis were performed as described by Suginta et al. [17, 12]. The lipid bilayer cuvette consisted of two chambers with a 25-μm thick Teflon film sandwiched in between. The latter had an aperture of 50-100 μm diameter, across which a virtually solvent-free planar lipid bilayer was formed. The chambers were filled with electrolyte solution and Ag/AgCl electrodes immersed on either side of the Teflon film. The electrolyte used was 1 M KCl buffered with 20 mM HEPES, pH 7.5. 1,2-Diphytanoyl-sn-glycero-3-phosphatidylcholine (DPhPC; Avanti Polar Lipids, Alabaster, AL) was used for lipid bilayer formation. To form the bilayer the aperture was first pre-painted with 1 μL of 1% (v/v) hexadecane in pentane (Sigma Aldrich). One of the electrodes was used as ground (*cis*) while the other electrode (*trans*) was connected to the headstage of an Axopatch 200B amplifier (Axon Instruments, Foster City, CA). Trimeric *Vh*ChiP channel (20-100 ng) was added to the solution on the *cis* side of the lipid membrane. At applied transmembrane potentials of ±100 mV, a single channel was frequently inserted within a few minutes. The protein solution in the chamber was gently diluted by multiple additions of the working electrolyte, to prevent multiple insertions. Single-channel current measurements were performed with an Axopatch 200B amplifier (Molecular Devices, Sunnywale., CA, U.S.A.) in the voltage-clamp mode, with the internal filter set at 10 kHz. Amplitude, probability and single-channel analyses were performed using pClamp v.10.5 software (all from Molecular Devices, Sunnyvale, CA). To investigate sugar translocation, chitooligosaccharide was added to the *cis* side of the chamber to a final concentration of 5 μM. Occlusions of ion flow caused by sugar diffusion through the inserting channel were usually recorded for 2 min. To determine the effect of sugar translocation on individual subunit blockages, discrete concentrations of chitohexaose (1.0, 2.5 and 5.0 μM) and chitotetraose (5.0, 10, 20 and 50 μM) were tested.

For the event detection of single-channel data, the ignore duration was set to 0.1 ms for chitohexaose and 0.05 ms for chitotetraose, to reject flickering and to avoid any collision peaks from the substrate. The kinetics of sugar binding to the channel were assessed by ion current fluctuation analysis [57]. For a given concentration [c] of chitooligosaccharide, the on-rate *k*_on_ (M^−1^·s^−1^) is obtained from the number of ion current blockage events (*v*) divided by the number of monomer closures in the trimeric form and the sugar concentration [*c*]:

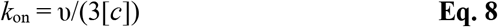

Reopening of the channel is a statistical event that is correlated to the strength of the binding. A channel closed at t = 0 has a probability *R*(t) of opening at time t. According to the simple binding model described above, an exponential function *R*(t) = *e*^−t/τ^ is expected. Fitting the lifetime distribution of a closed channel by an exponential parameter to obtain τ (the dwell time) yields the off-rate or *k*_off_ (s^−1^):

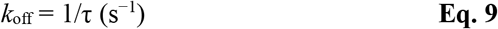

Alternatively, the equilibrium binding constant (*K*, M^−1^) was determined from a non-linear plot of conductance change versus sugar concentration using noise analysis as given in **Eq. 10** [59,60]:

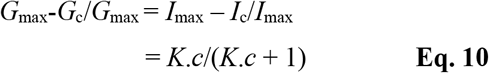

where *G*_max_ is the average conductance of the fully open *Vh*ChiP channel; *G*_c_ is the average conductance at a given chitooligosaccharide concentration; *I*_max_ is the initial current of the fully open channel without sugar; and *I*_c_ is the ionic current at a particular concentration of sugar.

## Abbreviations

BCA: bicinchoninic acid
ChiA: chitinase A
CD: circular dichroism
DNP-lysine: 2,4-dinitrophenyl-lysine
FPLC: fast protein liquid chromatography
D-GlcNAc: *N*-acetylglucosamine
(GlcNAc)_*n;*_ *n* = 2-6: chitooligosaccharides
octyl-POE: poly(ethylene glycol) octyl ether
LDAO: *N*,*N*-lauryldimethylamine oxide
*K*: equilibrium binding constant
*K*_d_: equilibrium dissociation constant
Omp: outer-membrane protein
SDS-PAGE: sodium dodecyl sulphate polyacrylamide gel electrophoresis
*T*_m_: midpoint temperature of thermal unfolding state
W136A: *Vh*ChiP W136A mutant
WT: *Vh*ChiP wild-type *Vh*ChiP

## Data availability

All data described in this publication will be shared upon request. Please contact.

## Acknowledgements

We would like to thank Dr. Takashi Ohhnuma, Kindai University, Japan for providing some of chitooligosaccharides. We thank Professor Mathias Winterhalter, Jacobs University Bremen, Bremen, Germany for introducing our lab members to single channel recordings using the BLM technique, with intensive training. CD measurements were carried out at the Salaya Equipment Central Equipment Facility, Mahidol University, Salaya Campus, Nakhon Prathom, Thailand. Finally, we would like to thank Dr. David Apps, the University of Edinburgh, U.K. for critical reading and English improvement of the manuscript. Special thanks to Professor Tamo Fukamizo from Kindai University, Japan for valuable discussion on cooperative characteristics of thermal unfolding by CD and fluorescence spectroscopy.

## Author contributions

AA carried out all experiments (except single channel electrophysiology of mutant W136A) and data analysis described in this study and took part in manuscript draft preparation. WCS prepared the first manuscript draft and took part in revisions. WC carried out single channel electrophysiology of *Vh*ChiP W136A mutant with chitohexaose and chitotetraose. AS provided technical guidance on electrophysiology and took part in manuscript draft preparation. WS conceived and initiated the project, supplied chemicals and materials, provided guidance throughout this study and wrote and revised the manuscript. All authors were informed and agree with the submission of the manuscript.

## Funding and additional information

AA was supported by full-time Postdoctoral scholarships from Suranaree University of Technology (Grant no Full-time61/04/2562) and the Postdoctoral Fellowship from Vidyasirimedhi Institute of Science and Technology. WS was funded by Start-up Grant from Vidyasirimedhi Institute of Science and Technology (Grant no: 300/111100/1711111000030) and the Thailand Research Fund through The Basic Research Grant (Grant no: BRG610008). WS also received financial support from Thailand Science Research and Innovation (TSRI) under the Global Partnership Grant.

## REFERENCES

1. Yan, N.; Chen, X., Sustainability: Don’t waste seafood waste. Nature 2015, 524 (7564), 155–157.

2. Arnold, N. D.; Brück, W. M.; Garbe, D.; Brück, T. B., Enzymatic modification of native chitin and conversion to specialty chemical products. Mar Drugs 2020, 18 (2), 93.

3. Yu, C.; Lee, A. M.; Bassler, B. L.; Roseman, S., Chitin utilization by marine bacteria. J Bio Chem 1991, 266 (36), 24260–24267.

4. Wang, S. L.; Chang, W. T.; Purification and characterization of two bifunctional chitinases/lysozymes extracellularly produced by *Pseudomonas aeruginosa* K-187 in a shrimp and crab shell powder medium. Appl Environ Microbiol 1997, 63 (2), 380–386.

5. Li, S. W.; He, H.; Zeng, R. J.; Sheng, G. P., Chitin degradation and electricity generation by *Aeromonas hydrophila* in microbial fuel cells. Chemosphere 2017, 168, 293–299.

6. Gurav, R.; Bhatia, S. K.; Choi, T. R.; Jung, H. R.; Yang, S. Y.; Song, H. S.; Park, Y. L.; Han, Y. H.; Park, J. Y.; Kim, Y. G.; Choi, K. Y.; Yang, Y. H., Chitin biomass powered microbial fuel cell for electricity production using halophilic *Bacillus circulans* BBL03 isolated from sea salt harvesting area. Bioelectrochem 2019, 130, 107329.

7. Sun, X.; Li, Y.; Tian, Z.; Qian, Y.; Zhang, H.; Wang, L., A novel thermostable chitinolytic machinery of Streptomyces sp. F-3 consisting of chitinases with different action modes. Biotechnol Biofuels 2019, 12 (1), 136.

8. Marmouzi, I.; Ezzat, S. M.; Salama, M. M.; Merghany, R. M.; Attar, A. M.; El-Desoky, A. M.; Mohamed, S. O., Recent Updates in Pharmacological Properties of Chitooligosaccharides. Biomed Res Int 2019, 2019, 4568039.

9. Kumar, M.; Brar, A.; Vivekanand, V.; Pareek, N., Bioconversion of Chitin to Bioactive Chitooligosaccharides: Amelioration and Coastal Pollution Reduction by Microbial Resources. Mar Biotechnol (NY) 2018, 20 (3), 269–281.

10. Ransangan, J.; Mustafa, S., Identification of *Vibrio harveyi* isolated from diseased Asian Seabass Lates calcarifer by use of 16S ribosomal DNA sequencing. J Aquat Anim Health 2009, 21 (3), 150–155.

11. Tendencia, E. A., *Vibrio harveyi* isolated from cage-cultured seabass Lates calcarifer Bloch in the Philippines. Aquacult Res 2002, 33 (6), 455–458.

12. Vezzulli, L.; Previati, M.; Pruzzo, C.; Marchese, A.; Bourne, D. G.; Cerrano, C., *Vibrio* infections triggering mass mortality events in a warming Mediterranean Sea. Environ Microbiol 2010, 12 (7), 2007–2019.

13. Lightner, D. V., Diseases of cultured penaeid shrimp. CRC handbook of mariculture 1993, 1, 393–486.

14. Suginta, W.; Robertson, P. A.; Austin, B.; Fry, S. C.; Fothergill-Gilmore, L. A., Chitinases from *Vibrio*: activity screening and purification of chiA from *Vibrio carchariae*. J Appl Microbiol 2000, 89 (1), 76–84.

15. Suginta, W.; Vongsuwan, A.; Songsiriritthigul, C.; Prinz, H., Estibeiro, P.; Duncan, R. R.; Svasti, J.; Fothergill-Gilmore, L. A., An endochitinase A from *Vibrio carchariae*: cloning, expression, mass and sequence analyses, and chitin hydrolysis. Arch Biochem Biophys 2004, 424 (2), 171–180.

16. Keyhani, N. O.; Li, X. B.; Roseman, S., Chitin catabolism in the marine bacterium *Vibrio furnissii*. Identification and molecular cloning of a chitoporin. J Biol Chem 2000, 275 (42), 33068–33076.

17. Suginta, W.; Chumjan, W.; Mahendran, K. R.; Janning, P.; Schulte, A.; Winterhalter, M., Molecular uptake of chitooligosaccharides through chitoporin from the marine bacterium *Vibrio harveyi*. PLoS One 2013, 8 (1), e55126.

18. Suginta, W.; Chuenark, D.; Mizuhara, M.; Fukamizo, T., Novel β-*N*-acetylglucosaminidases from *Vibrio harveyi* 650: cloning, expression, enzymatic properties, and subsite identification. BMC Biochem 2010, 11 (1), 40.

19. Jung, B. O.; Roseman, S.; Park, J. K., The central concept for chitin catabolic cascade in marine bacterium. Vibrios Macromol Res 2008, 16 (1), 1–5.

20. Hunt, D. E.; Gevers, D.; Vahora, N. M.; Polz, M. F., Conservation of the chitin utilization pathway in the *Vibrionaceae*. Appl Environ Microbiol 2008, 74 (1), 44–51.

21. Bassler, B. L.; Yu, C.; Lee, Y. C.; Roseman, S., Chitin utilization by marine bacteria: Degradation and catabolism of chitin oligosaccharides by *Vibrio furnissii*. J Biol Chem 1991, 266 (36), 24276–24286.

22. Li, X.; Roseman, S., The chitinolytic cascade in *Vibrios* is regulated by chitin oligosaccharides and a two-component chitin catabolic sensor/kinase. Proc Natl Acad Sci U S A 2004, 101 (2), 627–631.

23. Suginta, W.; Chumjan, W.; Mahendran, K. R.; Schulte, A.; Winterhalter, M., Chitoporin from *Vibrio harveyi*: a channel with exceptional sugar specificity. J Biol Chem 2013, 288 (16), 11038–11046.

24. Aunkham, A.; Zahn, M.; Kesireddy, A.; Pothula, K. R.; Schulte, A.; Baslé, A.; Kleinekathöfer, U.; Suginta, W.; van den Berg, B., Structural basis for chitin acquisition by marine *Vibrio* species. Nat Commun 2018, 9 (1), 1–13.

25. Soysa, H. S.; Suginta, W., Identification and Functional Characterization of a Novel OprD-like Chitin Uptake Channel in Non-chitinolytic Bacteria. J Biol Chem 2016, 291 (26), 13622–13633.

26. Corrêa, D.; Carlos, R., The use of circular dichroism spectroscopy to study protein folding, form and function. Afr J Biochem Res 2009, 3 (5), 164–173.

27. Miles, A. J.; Wallace, B. A., Circular dichroism spectroscopy of membrane proteins. Chem Soc Rev 2016, 45 (18), 4859–4872.

28. Greenfield, N. J., Using circular dichroism collected as a function of temperature to determine the thermodynamics of protein unfolding and binding interactions. Nat Protoc 2006, 1 (6), 2527–2535.

29. Schirmer, T.; Keller, T. A.; Wang, Y. F.; Rosenbusch, J. P., Structural basis for sugar translocation through maltoporin channels at 3.1 Å resolution. Science 1995, 267 (5197), 512–514.

30. Ishii, J. N.; Okajima, Y.; Nakae, T., Characterization of lamB protein from the outer membrane of Escherichia coli that forms diffusion pores selective for maltose-maltodextrins. FEBS Lett 1981, 134 (2), 217–220.

31. Forst, D.; Welte, W.; Wacker, T.; Diederichs, K., Structure of the sucrose-specific porin ScrY from *Salmonella typhimurium* and its complex with sucrose. Nat Struc Biol 1998, 5 (1), 37–46.

32. van den Berg, B.; Bhamidimarri, S. P.; Prajapati, J. D.; Kleinekathöfer, U.; Winterhalter, M., Outer-membrane translocation of bulky small molecules by passive diffusion. Proc Natl Acad Sci U S A 2015, 112 (23), E2991–E2999.

33. van den Berg, B., Structural basis for outer membrane sugar uptake in *Pseudomonads*. J Biol Chem 2012, 287 (49), 41044–41052.

34. Biswas, S.; Mohammad, M. M.; Patel, D. R.; Movileanu, L.; van den Berg, B., Structural insight into OprD substrate specificity. Nat Struct Mol Biol 2007, 14 (11), 1108–1109.

35. Suginta, W.; Smith, M. F., Single-molecule trapping dynamics of sugar-uptake channels in marine bacteria. Phys Rev Lett 2013, 110 (23), 238102.

36. Suginta, W.; Winterhalter, M.; Smith, M. F., Correlated trapping of sugar molecules by the trimeric protein channel chitoporin. Biochim Biophys Acta 2016, 1858 (12), 3032–3040.

37. Takeuchi, T.; Morita, K.; Saito, T.; Kugimiya, W.; Fukamizo, T., Chitosan-soyprotein interaction as determined by thermal unfolding experiments. Biosci Biotechnol Biochem 2006, 70 (7), 1786–1789.

38. Honda, Y.; Fukamizo, T.; Okajima, T.; Goto, S.; Boucher, I.; Brzezinski, R., Thermal unfolding of chitosanase from *Streptomyces sp*. N174: role of tryptophan residues in the protein structure stabilization. Biochimica et Biophysica Acta 1999, 1429 (2), 365–376.

39. Honda, Y.; Fukamizo, T.; Okajima, T.; Goto, S.; Boucher, I.; Brzezinski, R., Thermal unfolding of chitosanase from Streptomyces sp. N174: role of tryptophan residues in the protein structure stabilization. Biochim Biophys Acta 1999, 1429 (2), 365–376.

40. Fukamizo, T.; Juffer, A. H.; Vogel, H. J.; Honda, Y.; Tremblay, H.; Boucher, I.; Neugebauer, W. A.; Brzezinski, R., Theoretical calculation of *p*Ka reveals an important role of Arg205 in the activity and stability of Streptomyces sp. N174 chitosanase. J Biol Chem 2000, 275 (33), 25633–25640.

41. Dutzler, R.; Schirmer, T.; Karplus, M.; Fischer, S., Translocation mechanism of long sugar chains across the maltoporin membrane channel. Structure 2002, 10 (9), 1273–1284.

42. Forst, D.; Welte, W.; Wacker, T.; Diederichs, K., Structure of the sucrose-specific porin ScrY from *Salmonella typhimurium* and its complex with sucrose. Nat Struct Biol 1998, 5 (1), 37–46.

43. Ross, P. D.; Subramanian, S., Thermodynamics of protein association reaction: forces contribution to stability. Biochemistry 1981, 20 (11), 3096–3102.

44. Wallace, A. C.; Laskowski, R. A.; Thornton, J. M., LIGPLOT: a program to generate schematic diagrams of protein-ligand interactions. Protein Eng 1995, 8 (2), 127–134.

45. Chumjan, W.; Winterhalter, M.; Schulte, A.; Benz, R.; Suginta, W., Chitoporin from the Marine Bacterium *Vibrio harveyi*: Probing the essential roles of Trp136 at the surface of the construction zone. J Biol Chem 2015, 290 (31), 19184–19196.

46. Prilipov, A.; Phale, P. S.; Van Gelder, P.; Rosenbusch, J. P.; Koebnik, R., Coupling site-directed mutagenesis with high-level expression: large scale production of mutant porins from *E. coli*. FEMS Microbiol Lett 1998, 163 (1), 65–72.

47. Aunkham, A.; Schulte, A.; Winterhalter, M.; Suginta, W., Porin involvement in cephalosporin and carbapenem resistance of *Burkholderia pseudomallei*, PLoS One 2014, 9 (5), e95918.

48. Rosenbusch, J. P., Characterization of the major envelope protein from *Escherichia coli*. Regular arrangement on the peptidoglycan and unusual dodecyl sulfate binding. J Biol Chem 1974, 249 (24), 8019–8029.

49. Lugtenberg, B.; van Alphen, L., Molecular architecture and functioning of the outer membrane of *Escherichia coli* and other Gram-negative bacteria. Biochim Biophys Acta 1983, 737 (1), 51–115.

50. Mallik, B.; Chakravarti, B.; Chakravarti, D. N., Overview of Chromatography. Current Protocols Essential Laboratory Techniques 2008, (1), 6–1.

51. Riley, M. L.; Wallace, B. A.; Flitsch, S. L.; Booth, P. J., Slow alpha helix formation during folding of a membrane protein. Biochemistry 1997, 36 (1), 192–196.

52. Sehgal, P.; Otzen, D. E., Thermodynamics of unfolding of an integral membrane protein in mixed micelles. Protein Sci 2006, 15 (4), 890–899.

53. Rinnenthal, J.; Klinkert, B.; Narberhaus, F.; Schwalbe, H., Modulation of the stability of the Salmonella fourU-type RNA thermometer. Nucleic Acids Res 2011, 39 (18), 8258–8270.

54. Songsiriritthigul, C.; Pantoom, S.; Aguda, A. H.; Robinson, R. C.; Suginta, W., Crystal structures of *Vibrio harveyi* chitinase A complexed with chitooligosaccharides: implications for the catalytic mechanism. J Struct Biol 2008, 162 (3), 491–499.

55. Srivastava, D. B.; Ethayathulla, A. S.; Kumar, J.; Singh, N.; Sharma, S.; Das, U.; Srinivasan, A.; Singh, T. R., Crystal structure of a secretory signaling glycoprotein from sheep at 2.0 Å resolution. J Struct Biol 2006, 156 (3), 505–516.

56. Neves, P.; Berkane, E.; Gameiro, P.; Winterhalter, M.; Castro, B., Interaction of quinolones antibiotics and bacterial outer membrane porin OmpF. Biophys Chem 2005, 113 (2), 123–128.

57. Kullman, L.; Winterhalter, M.; Bezrukov, S. M., Transport of maltodextrins through maltoporin: a single-channel study. Biophys J 2002, 82 (2), 803–812.

58. Sun, S. F.; Zhou, B.; Hou, H. N.; Liu, Y.; Xiang, G. Y., Studies on the interaction between Oxaprozin-E and bovine serum albumin by spectroscopic methods. Int J Biol Macromol 2006, 39 (4-5), 197–200.

59. Benz, R.; Hancock, R. E., Mechanism of ion transport through the anion-selective channel of the *Pseudomonas aeruginosa* outer membrane. J Gen Physiol 1987, 89 (2), 275–295.

60. Andersen, C.; Jordy, M.; Benz, R., Evaluation of the rate constants of sugar transport through maltoporin (LamB) of *E. coli* from the sugar-induced current noise. J Gen Physiol 1995, 105 (3), 385–401.

